# A gene regulatory network critical for axillary bud dormancy directly controlled by Arabidopsis BRANCHED1

**DOI:** 10.1101/2020.12.14.394403

**Authors:** Sam W. van Es, Aitor Muñoz-Gasca, Francisco J. Romero-Campero, Eduardo González-Grandío, Pedro de los Reyes, Carlos Tarancón, Aalt D.J. van Dijk, Wilma van Esse, Gerco C. Angenent, Richard Immink, Pilar Cubas

## Abstract

The control of branch outgrowth is critical for plant fitness, stress resilience and crop yield. The *Arabidopsis thaliana* transcription factor BRANCHED1 (BRC1) plays a pivotal role in this process as it integrates signals that inhibit axillary bud growth to control shoot branching. Despite the remarkable activity of BRC1 as a potent growth inhibitor, the mechanisms by which it promotes and maintains bud dormancy are still largely unknown.

Here we combine ChIP-seq, transcriptomic and systems biology approaches to characterise the BRC1-regulated gene network. We identify a group of BRC1 direct target genes encoding transcription factors (BTFs) that orchestrate, together with BRC1, an intricate transcriptional network enriched in abscisic acid signalling components. The BRC1 network is enriched in feed-forward loops and feed-back loops, robust against noise and mutation, reversible in response to stimuli, and stable once established. This knowledge is fundamental to adapt plant architecture and crop production to ever-changing environmental conditions.

## Introduction

In multicellular organisms, growth and development occur by the sequential activation of gene regulatory networks (GRNs) that influence patterns of cell division expansion and differentiation in developing tissues, which results in species-specific morphologies. Sessile organisms, such as plants, need to adapt these developmental programs to resource availability and environmental conditions for fitness and survival. Thus, the activity of these GRNs, controlled by transcriptional master regulators, is modulated by external signals, such as light quality, photoperiod and temperature, and endogenous cues that signal nutrient and energy limitations or stress.

In higher plants, one key developmental program that determines aboveground plant architecture is the formation of branches, which are lateral shoots derived from axillary meristems. Axillary meristems are groups of undifferentiated cells formed in the leaf axils, which generate axillary buds or branch primordia. Under favourable conditions, these axillary buds continue to grow and elongate to form branches. In contrast, under actual or potentially energy-limiting situations, the buds remain dormant^1^. The status of each bud, either active or quiescent, is thus influenced by the environment and coordinated at the plant level by sugar availability, strigolactone (SL) hormone signalling, and auxin transport^2^. However, the molecular operations required for cells in axillary buds to integrate those signals locally, and implement the pathways of dormancy or growth are still poorly understood.

In *Arabidopsis thaliana, BRANCHED1* (*BRC1*) is a central local regulator of bud activity that encodes a <u>T</u>EOSINTE BRANCHED1, <u>C</u>YCLOIDEA, <u>P</u>CF (TCP) ^3^transcription factor (TF), whose function is widely conserved in angiosperms. *BRC1* and its orthologs are expressed in axillary buds and promote bud growth arrest: *brc1* mutants have increased axillary bud activity and excessive branching in a wide variety of species^2^. Due to the spatially restricted expression patterns of *BRC1*, limited to axillary buds, this gene only prevents lateral shoot outgrowth. However, when ectopically expressed in Arabidopsis seedlings, it can also cause growth arrest of shoot and root apical meristems and leaf primordia^4^. In hybrid aspen, *BRC1-like* genes expressed in the shoot apex in short days also promote growth cessation^5^. Despite these remarkable growth-inhibiting effects of *BRC1*, and its major role in the control of plant architecture, the gene targets and GRNs controlled directly by BRC1 remain largely unknown in Arabidopsis. Three closely related HD-zip-I genes (*HB21, HB40*, and *HB53*) are direct BRC1 targets and mediate *BRC1*-induced abscisic acid (ABA) synthesis and response in buds. ABA signalling in turn promotes inhibition of bud growth^6^. Furthermore, two clusters of co-expressed genes -one enriched in genes related to cell division and DNA replication, another in chloroplast and ribosomal genes-are downregulated in response to *BRC1*, although it is unclear whether this control is direct^4^. In maize, the closely related TCP factor Teosinte Branched1 (TB1) directly influences hormone signalling (ABA, gibberellic acid (GA), and jasmonate (JA)) and sugar signalling^7^ but it is unclear whether the TB1 targets are shared with Arabidopsis BRC1, as there is abundant evidence of the divergent regulatory pathways associated with TB1 and BRC1.

In this work, we have characterised the transcriptional network that operates directly downstream of Arabidopsis BRC1 to promote axillary bud growth arrest. For this, we have compared the transcriptome of seedlings after *BRC1* induction, the genome-wide binding sites of BRC1 by Chromatin Immunoprecipitation sequencing (ChIP-seq), and the transcriptional profiling of active vs dormant axillary buds. We have identified a group of direct targets of BRC1 encoding TFs that we have termed BTFs, which play a critical role in the regulation of the network. We have integrated this BRC1 ChIP-seq data with ChIP-/DAP-seq data of the *BRC1*-dependent TFs into a regulatory network, BRC1NET. To better explore and characterise the network properties we have developed an open interactive web application. Our findings reveal a pivotal role of BRC1 as a master regulator of an intricate transcriptional network enriched in ABA-related components. This transcriptional network, robust against noise and mutation, is reversible in the initial steps but becomes stable once established. These properties help fine-tune the transcriptional responses promoting bud dormancy in response to limiting conditions, an adaptation critical for plant fitness and survival.

## Results

### *BRC1* regulates nine coexpressed gene clusters

To identify the genes regulated by BRC1, our first approach was to perform a thorough *in silico* analysis of existing expression data. González-Grandío et al. (2013) carried out transcriptional profiling of axillary buds of wild-type and *brc1* mutant plants exposed for eight hours to either white light (WL), or white light enriched in far-red light (WL+FR). The latter treatment simulates a canopy shade, which leads to *BRC1* mRNA accumulation and promotes bud dormancy. We used the collection of genes differentially expressed (DE, FDR<0.05) in response to *BRC1* (*BRC1*-dependent genes, **Supplemental Figure 1**) as a starting point for this study. To investigate their potential functional relationships, we studied their degree of co-expression in > 15,200 publicly available microarray experiments and clustered them by their global similarity of transcriptional responses. This analysis revealed six co-expression clusters of up-regulated genes and three co-expression clusters of down-regulated genes (**Fig. 1a**). We then searched *in silico* for additional genes co-expressed with members of each cluster and tested whether they were also DE in dormant (WL+FR-treated wild-type) buds, but not in active (WL-treated wild-type, or *brc1* mutant) buds (**Fig. 1b**). Genes that met this criterion were added to the *BRC1*-dependent category (**Supplemental Fig. 1 and 2)**. We termed these clusters of co-expressed genes, **UP_C1** to **C6** and **DOWN_C1** to **C3,** for the up- and down-regulated genes in dormant buds, respectively (**Fig. 1b, Supplemental Dataset 1**).

**Fig. 1.**
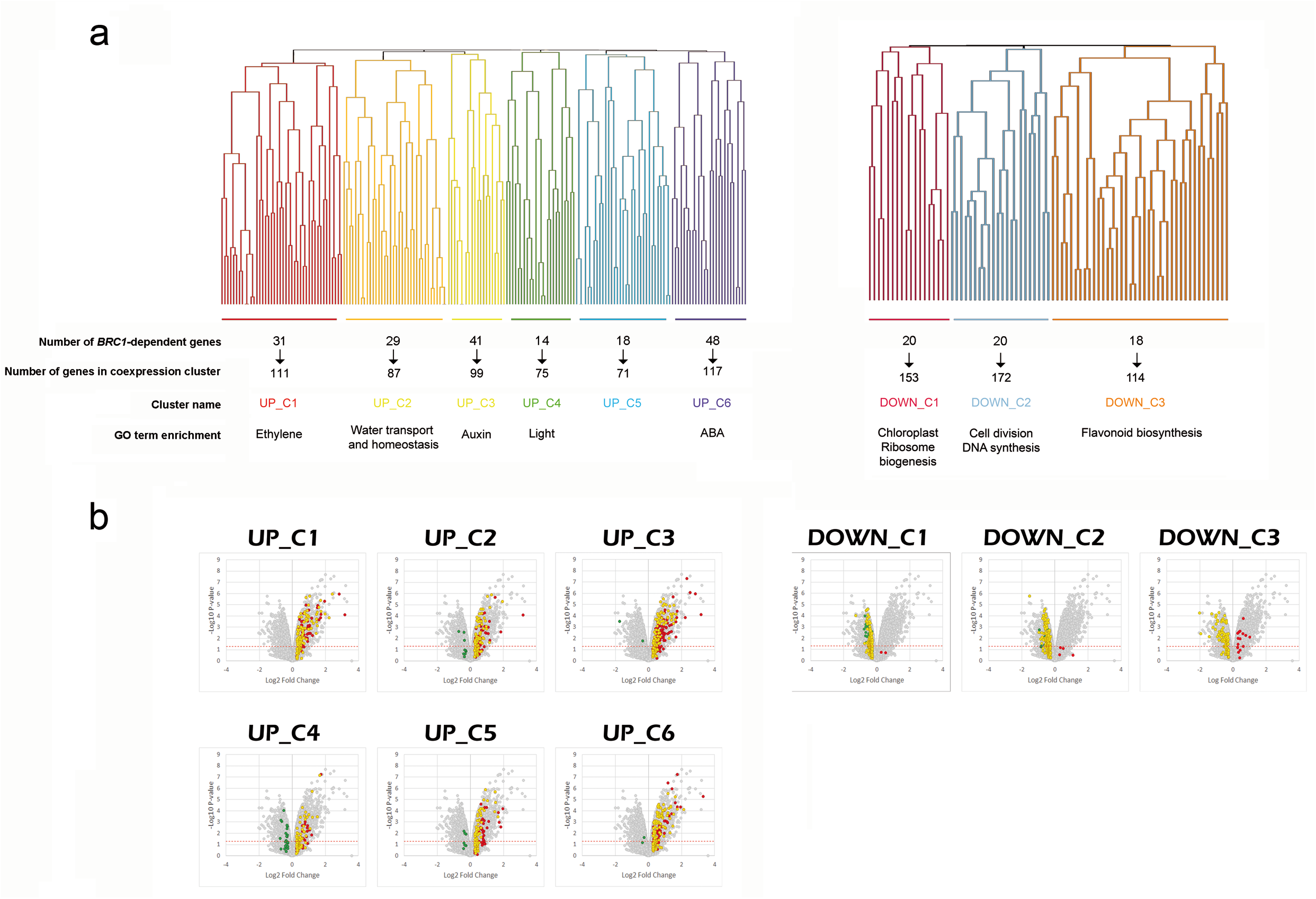
*BRC1*-dependent gene coexpression clusters. **a**, Hierarchical clustering of *BRC1*-dependent genes^4^ based on their degree of coregulation in 15,275 microarray experiments (ATTED-II^36^). Only 238 (181 UP and 57 DOWN) of the 307 genes are represented in Affimetrix arrays and could be analysed. Number of coregulated genes, number of additional co-expressed genes, cluster name and overrepresented GO terms are indicated for each cluster. **b**, Validation of genes co-expressed with BRC1 dependent genes. Volcano plots representing pval (−Log10 pval, vertical axis) and relative expression (Log2 fold change, horizontal axis) of genes in the active vs dormant buds (8 h low R:FR vs. High R:FR) microarray. *BRC1*-dependent genes and their co-expressed genes are highlighted. Genes highlighted in yellow are the original *BRC1* dependent genes plus those co-expressed genes induced or repressed only in wild type but not in *brc1* mutants. These genes were further analysed. In red, genes induced both in wild-type and *brc1* mutant buds. In green, genes repressed both in wild-type and *brc1* mutant buds. Green and red genes were not included in subsequent analyses.

Each co-expression cluster was enriched in a different set of gene ontology (GO) terms^9^ (**Fig. 1a, Supplemental Dataset 1**): **UP_C1** was enriched in genes involved in ethylene signalling, senescence, and catabolism; **UP_C2**, in genes related to water transport; **UP_C3**, brassinosteroid and auxin signalling; **UP_C4**, response to light; **UP_C5** did not show an statistically significant enrichment in GO terms, but contained genes related to cell wall remodelling and GA signalling, among others; **UP_C6**, was enriched in genes related to ABA signalling and response to abiotic stress; **DOWN_C1,** in genes related to chloroplast activity and organization; **DOWN_C2**, in DNA replication and cell division; and **DOWN_C3** in genes related to biosynthesis of secondary metabolites such as flavonoids, glucosinolates and pectins.

This collection of 947 genes constitutes a core transcriptional response associated with *BRC1* activity and to the growth-to-dormancy transition in Arabidopsis axillary buds.

### Identification of BRC1 direct targets

We next investigated how BRC1 orchestrates the expression of these *BRC1*-dependent gene clusters in buds. BRC1 could directly regulate the expression of a majority of genes from each cluster, or it could control a limited number of key regulators, which in turn would mediate a wider transcriptional response. To discern between these possibilities, we looked for direct BRC1 targets in the Arabidopsis genome using ChIP-seq^9^. *BRC1* expression levels are low and limited to axillary buds^10^, which complicate the study of protein-DNA binding events^11^. We therefore performed ChIP in ten-day-old seedlings of the estradiol-inducible line *LEXA*:*GFP*:*BRC1^ind^*; *brc1-2* (*BRC1^ind^*)^6^. We collected material five hours after the beginning of estradiol induction, when the GFP:BRC1 protein was detectable^6^. ChIP-seq was performed for three independent biological replicates (**Supplemental Fig. 3, Supplemental Dataset 2)**. We found that 4339 loci contained BRC1 peaks within 3 kb upstream of the TSS. A large proportion (~60%) of the BRC1 peaks were located in the proximal upstream region of protein-coding genes, on average within 300 bp of the transcription start site (TSS) (**Fig. 2a**). Around 4000 loci (~60%) contained peaks within 3 kb upstream of their TSS in the two replicates with the highest sequencing and mapping depth. The other peaks were positioned in introns, exons, or UTRs (totally ~ 9%), and the remaining ~10 and ~20% in the downstream and intergenic regions, respectively (**Fig. 2b)**. This distribution was comparable to that found for ChIP-seq performed on other Arabidopsis TFs (e.g.^12,13^), but divergent from that described for cucumber TEN, a closely related TCP factor, for which a large proportion of binding sites are localised in gene bodies^14^.

**Fig. 2.**
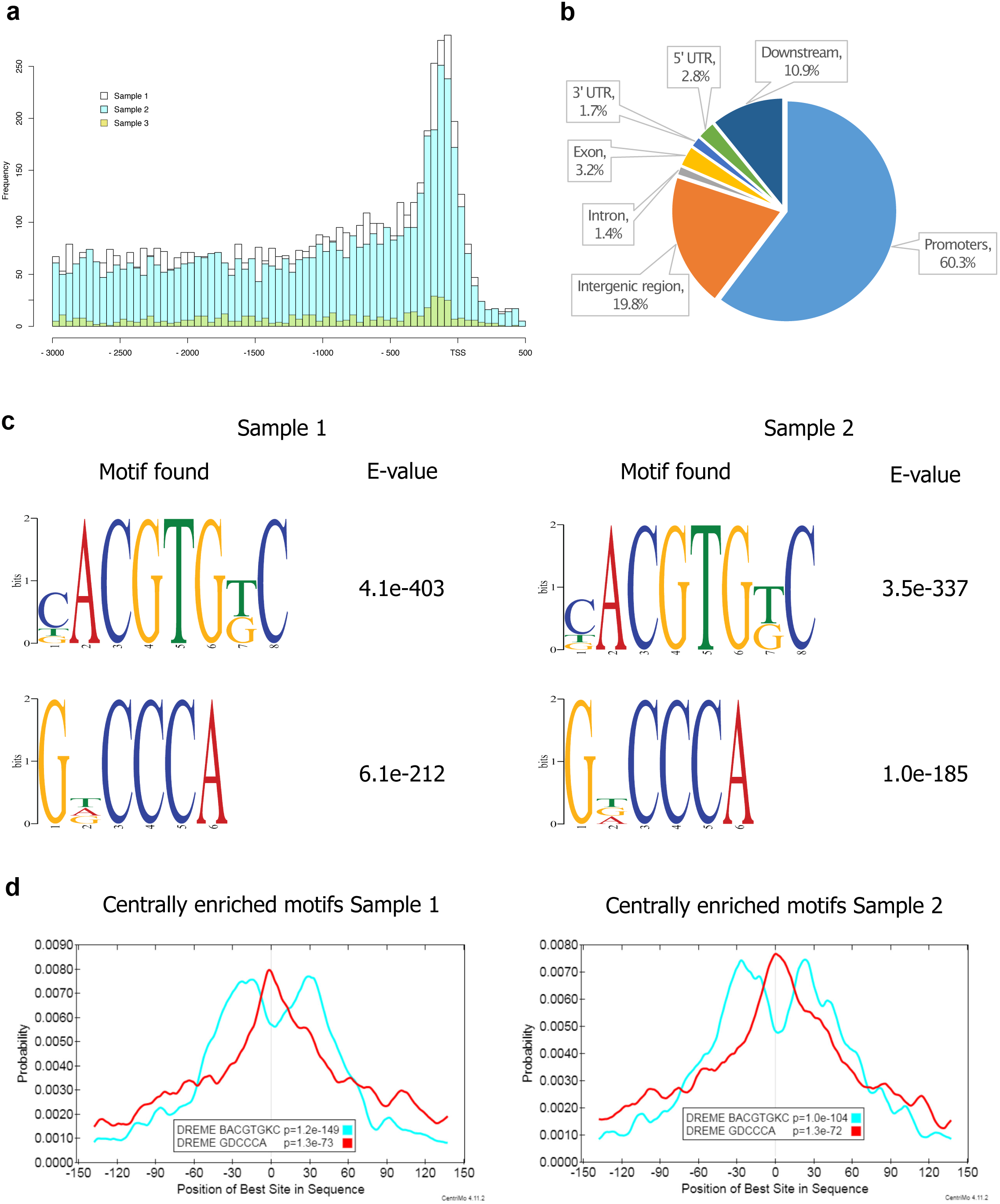
Overview of ChIP-seq data and MEME motif enrichment analysis. **a**, Sequencing read distribution 3kb up- to 500bp downstream of the transcription start site (TSS) in BRC1 bound loci. **b**, Distribution of reads over the different regions in the identified BRC1 bound loci (promoter, intron, exon etc.). **c**, Results of the MEME motif enrichment analysis. Specific enrichment was found for a consensus BRC1 TCP-like binding motif and a G-box motif. D. Localization of the identified motifs in the BRC1 bound regions. The GDCCCA (motif 2; TCP-binding motif) is centrally enriched, whereas the CACGTG (motif 1; G-box) is found enriched in short distance up- or downstream of the ChIP peak center.

BRC1-bound peaks in the two most enriched ChIP-seq samples showed a significant overrepresentation of several motifs (**Supplemental Table 1,** DREME^15^ most notably GDCCCA and BACGTGKC (D=AGT; B=CGT; K=GT)(**Fig. 2c**). Around 60% (57.5 and 65.1% for samples 1 and 2, respectively) of all peaks contained the GDCCCA motif, which closely resembles a TCP binding site^16,17^ and the BRC1 binding motif <u>GGgcCCmc</u>, inferred by Protein-binding microarray assays^6^. Remarkably, 50% of all peaks (48.3 and 51.9% for samples 1 and 2, respectively) contained the BACGTGKC motif, which closely resembles a G-box (CACGTG) recognised by bHLH and bZIP TFs^18^. Interestingly, in approximately 30% of the peaks (28.5 and 34.5% for samples 1 and 2, respectively), the TCP binding site was accompanied by a G-box-like motif. In these cases, the TCP motif was centrally enriched, whereas the G-box was positioned either up- or downstream of the TCP binding site (**Fig. 2d)**. This suggests that BRC1 frequently acts in concert with G-box binding proteins to control gene expression. The overrepresentation of a G-box-like motif was not reported in the recently published ChIP-seq experiment for maize TB1^7^ and this prompted us to reanalyse that dataset. Our analysis confirmed that the G-box is also overrepresented in the maize TB1 targets (**Supplemental Results**), revealing that this feature is indeed conserved in maize, a monocot distantly related to Arabidopsis.

In parallel to the ChIP study, we did a gene expression analysis (RNA-seq) in Arabidopsis of similar estradiol-treated material, to identify genes directly responding to *BRC1* activity (differentially expressed genes, DEG, **Supplemental Dataset 3**). We found 5718 DEGs of which 1845 were upregulated and 3873 were downregulated after induction (false discovery rate, FDR <0.05). We considered *BRC1 direct targets* those genes whose regulatory regions were bound by BRC1 (i.e. *BRC1-bound genes*) and were DE after *BRC1* induction in seedlings (**Supplemental Fig. 1**). We identified 1438 BRC1 direct targets according to this criterion.

### *BRC1*-dependent direct gene targets

To further refine the BRC1 target gene list, we compared the BRC1 direct targets identified by ChIP-seq and RNA-seq with the *BRC1*-dependent genes identified in the active-vs-dormant bud transcriptomics. Genes present in the three lists were termed *bona fide BRC1 direct targets* (**Supplementary Fig. 1, Supplemental Dataset 4**). We found 97 genes; 90 induced and 7 repressed in response to *BRC1*, both in seedlings and in buds entering dormancy. This strongly indicates that BRC1 acts mainly as a transcriptional activator. The list was enriched in genes containing ABA-related terms (ABA signalling, FDR= 0.045; ABA response, FDR= 7.4 10^×4^; ABA transport, FDR=0.037), and genes responding to water deprivation and salt/osmotic stress (FDR=0.01) (**Supplemental Dataset 4**). Another significantly enriched category (FDR=0.037) was that of regulation of transcription: 19 *bona fide* BRC1 direct targets encoded TFs, many of which had ABA-related annotations (**Table 1**). The list of *bona fide* BRC1 targets also contained induced genes encoding enzymes controlling hormone metabolism and homeostasis such as *MAX1, NCED3* and *ST2A* (that regulate SL, ABA, and JA synthesis respectively,^19–21^) Beta-carotene 3-hydroxylase 2 (*BCH2*, which catalyses the synthesis of zeaxanthin, a precursor of SL and ABA); *CYP707A2* and *GA2OX2* (involved in ABA and GA degradation respectively); and *WES1/GH3.5* (that control auxin amino acid conjugation). Among the 6 downregulated genes was *UGT76C2* (that promotes cytokinin glycosylation and inactivation). Direct targets of BRC1 are also genes encoding ubiquitination-related proteins, as well as water, ion, amino acid and hormone membrane transporters.

**Table 1.** BRC1-regulated Transcription factors (BTFs). List of *bona fide* BRC1 direct targets encoding TFs. In bold, those related to ABA signaling and/or response.

| AGI | Name | Description | TF family |
| --- | --- | --- | --- |
| **At4g36740** | **ATHB-40** | **Homeobox-leucine zipper protein ATHB-40** | **HD-zip** |
| **At5g66700** | **ATHB-53** | **Homeobox-leucine zipper protein ATHB-53** | **HD-zip** |
| **At2g18550** | **ATHB-21** | **Homeobox-leucine zipper protein ATHB-21** | **HD-zip** |
| **At4g01120** | **GBF2** | **G-box-binding factor 2** | **bZIP** |
| **At2g46270** | **GBF3** | **G-box-binding factor 3** | **bZIP** |
| **At4g34000** | **ABF3** | **ABA Responsive Element Binding Factor 3** | **bZIP** |
| **At2g36270** | **ABI5** | **ABA Insensitive 5** | **bZIP** |
| **At1g77450** | **ANAC32** | **NAC domain containing protein 32** | **NAC** |
| **At1g01720** | **ATAF1/ANAC002** | **ATAF1/ANAC domain-containing protein 2** | **NAC** |
| **At1g70000** | **MYBD** | **MYB-like transcription factor** | **MYB** |
| At2g40970 | MYBC1 | Homeodomain-like superfamily protein | Homeodomain |
| **At5g13330** | **ERF113/Rap2.6L** | **Ethylene-responsive transcription factor ERF113** | **ERF/AP2** |
| At3g61630 | CRF6 | Ethylene-responsive transcription factor CRF6 | ERF/AP2 |
| At5g49700 | AHL17 | AT-hook Motif Nuclear Localized Protein 17 | AT-hook |
| **At5g44260** | **TZF5** | **Tandem CCCH Zinc Finger protein 5** | **CCCH Zinc Finger** |
| AT5G62020 | HSFB2A/HSF6 | Heat Stress transcription factor B-2A | HSF |
| At1g19050 | ARR7 | Response Regulator 7 | ARR |
| **At4g31800** | **WRKY18** | **WRKY transcription factor 18** | **WRKY** |
| **At1g07090** | **LSH6** | **Light-dependent Short Hypocotyl 6** | **ALOG** |

A comparison of the ChIP-seq and RNA-seq data with the *BRC1*-dependent clusters showed that all the **UP** clusters but especially **UP_C4** and **UP_C6** were enriched in BRC1-bound, DEG and BRC1 direct targets (**Supplemental Fig. 4a,b,d**) whereas the **DOWN** clusters were not (**Supplemental Fig. 4a,c,e**).

All these results indicate that BRC1 controls directly the expression of numerous genes related to ABA function, and the activity and homeostasis of other hormones involved in bud activity such as SL, GA, auxin, and cytokinin^22^, as well as JA, which can effectively repress cell division^23^. BRC1 may also modulate the transport of nutrients, signals and water in and out of the axillary bud. Finally, BRC1 regulates a remarkable collection of TFs, which probably helps control the BRC1 transcriptional network.

### Some BRC1 direct target genes encode TFs that bind G-box motifs

*Bona fide* BRC1 direct targets were enriched in genes encoding TFs that we have termed BRC1-targeted TFs (BTFs): HD-Zip genes *HB21, HB40* and *HB53*; bZIP genes *GBF2, GBF3, ABF3*, and *ABI5*; NAC genes *ATAF1* and *NAC032*; AP2/EREB genes *ERF113/Rap2.6L* and *CRF6*; MYB genes *MYBC1*, and *MYBD*; zinc finger protein gene *ATTZF5*; and heat shock protein gene *HSB2A* (**Fig. 3, Table 1)**.

**Fig. 3.**
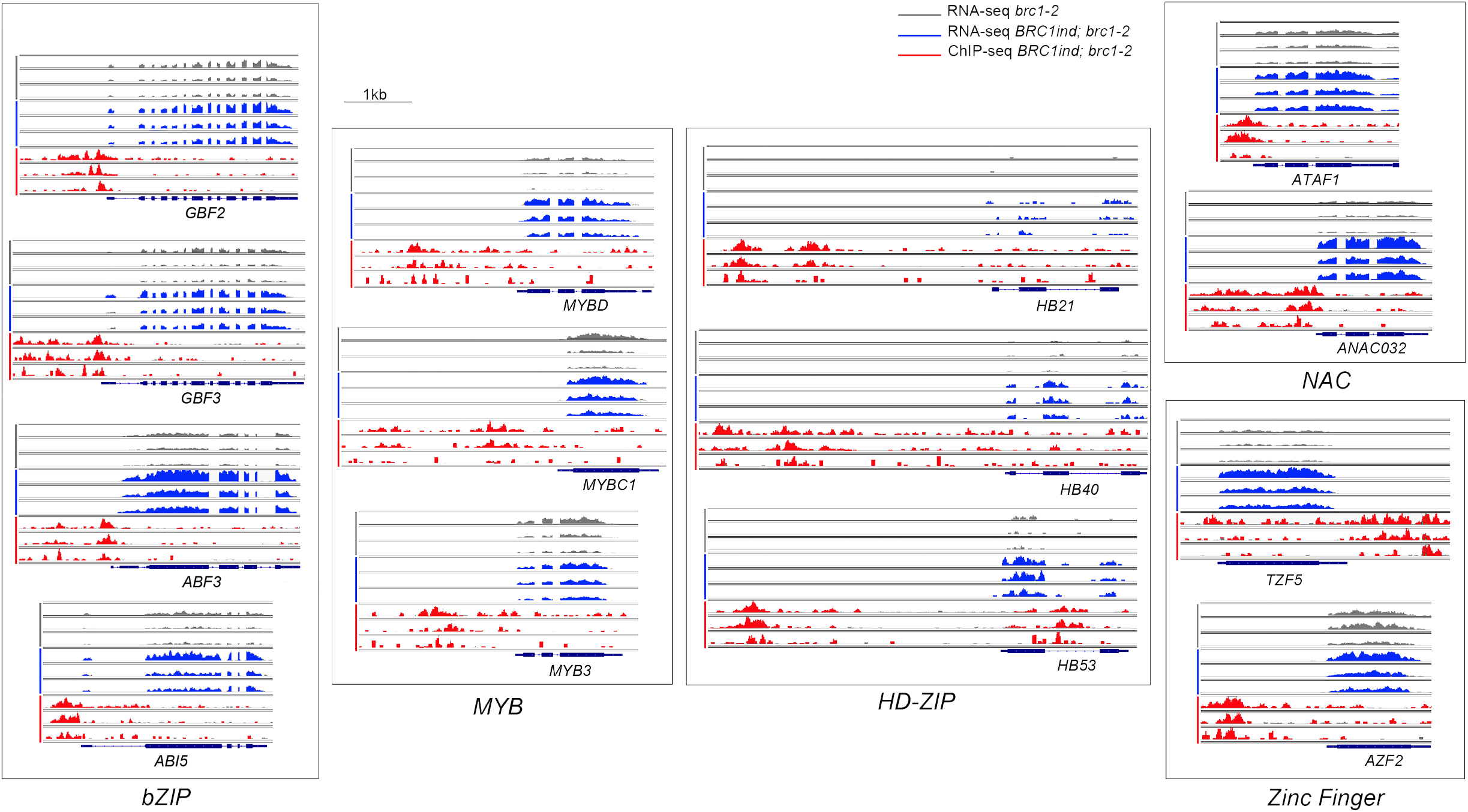
Transcription factors directly regulated by BRC1. BRC1 binding peaks (red) and *BRC1*-induced differential gene expression (blue) for 12 genes encoding BRC1 - regulated Transcription factors (BTFs) of the bZIP, NAC, MYB and Zinc Finger families. *MYB3* and *AZF2* (*BRC1*-dependent and BRC1-bound but not differentially expressed in seedlings) are also included as potential additional BRC1 targets in buds.

The promoters of some of these BTFs (*ABI5, ABF3, GBF2, GBF3, ATAF1*) were fused to the reporter β-*GLUCURONIDASE* and transformed into Arabidopsis. The resulting lines confirmed that, like *BRC1*^10^, these genes were strongly and specifically expressed in axillary buds (**Fig. 4a-i**). *HB21*, *HB40* and *HB53* are also expressed in these primordia^6^. Other *BRC1*-dependent, BRC1-bound genes encoding TFs (e.g. *NAP, MYB3*) were also highly expressed in axillary buds (**Fig. 5j-l**).

**Fig. 4.**
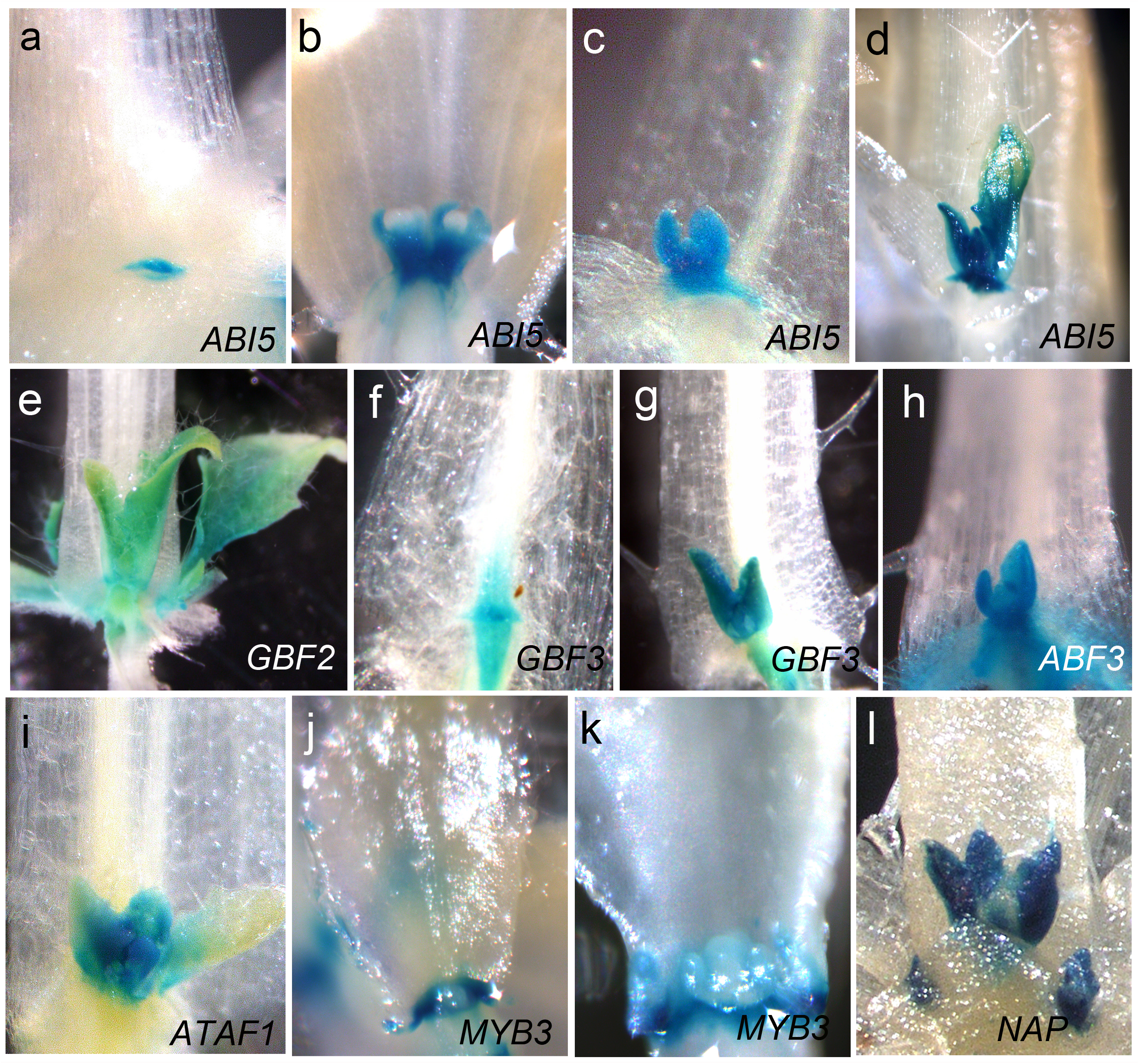
The BTFs are expressed in axillary buds. *GUS* activity in axillary buds of transgenic lines *ABI5p:GUS* (**a-d**), *GBF2p:GUS* (**e**), *GBF3p:GUS* (**f,g**), *ABF3p:GUS* (**h**), *ATAF1p:GUS* (**i**). The expression patterns of different BTFs are not identical: only *GBF3* and *ABF3* are expressed in the subtending vasculature. The expression of *MYB3* and *NAP*, BRC1-bound & *BRC1*-dependent genes, is also restricted to axillary buds as confirmed in *MYB3p:GUS* (**j, k**) and *NAPp:GUS* (**l**) transgenic lines. *MYB3* expression occurs at the base of the bud and is excluded from axillary meristems.

**Fig. 5.**
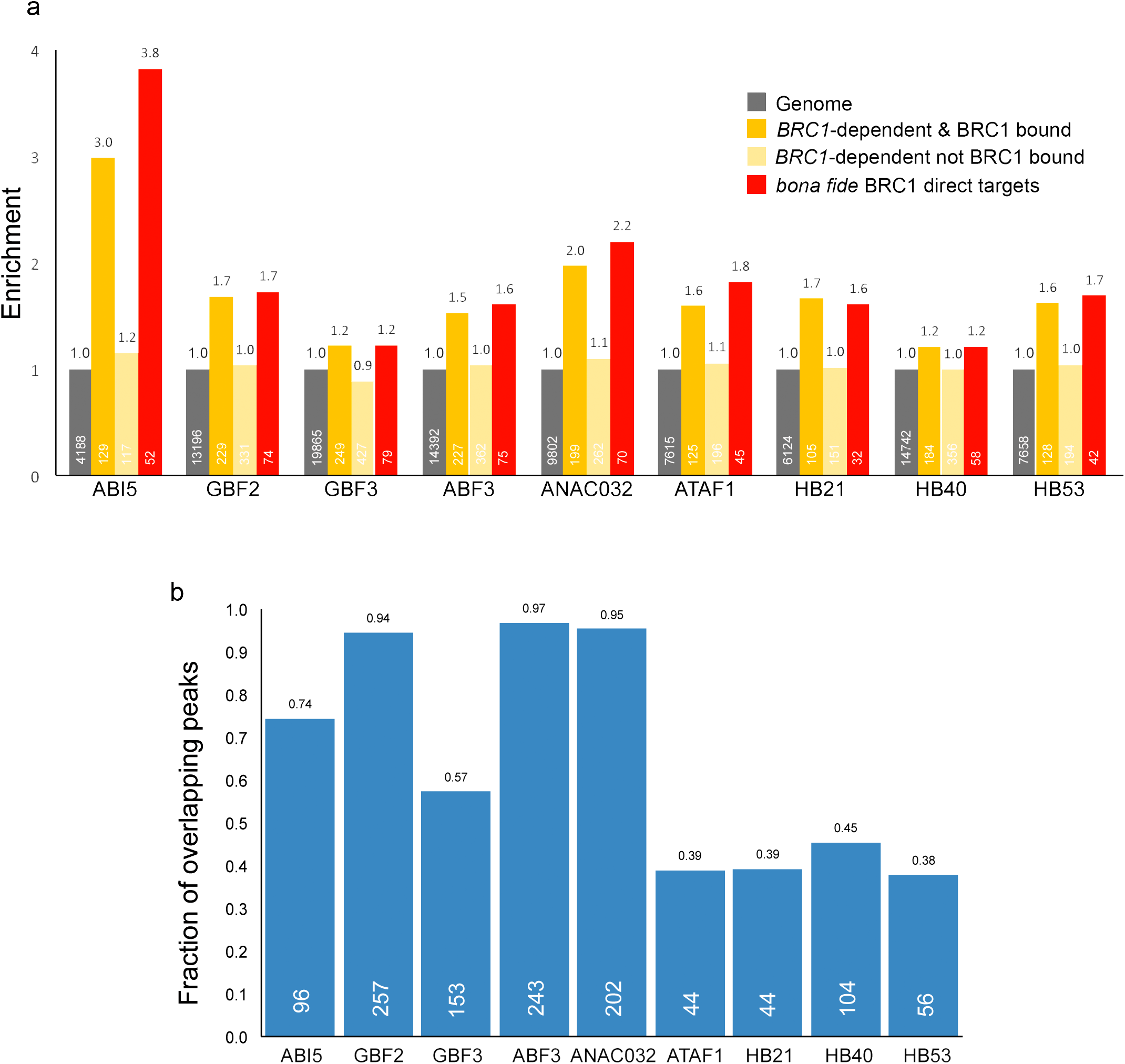
BRC1 and BTF binding sites largely overlap. **a**, Enrichment of BTF gene targets in the following gene sets: *BRC1*-dependent & BRC1-bound, *BRC1*-dependent not BRC1-bound and *bona fide* BRC1 direct targets. Numbers inside the bar indicate number of genes in each category. **b**, Fraction of BRC1 peaks found in BTF target gene promoters, which overlap with BTF peaks. Number inside bar indicates number of BRC1 peaks occurring in promoters of BTF target genes that overlap with BTF peaks.

Several of these BTFs (i.e. GBF2, GBF3, ABF3, ABI5, ANAC032, ATAF1) bind G-box-like motifs, in which BRC1 peaks are enriched (see above), and GBF2 and GBF3 also bind TCP-like motifs after ABA treatments (**Supplemental Fig. 5**^24^). This raises the possibility that BTFs act as co-regulators with BRC1 of the *BRC1*-dependent network. Consistently, *in silico* analyses in search for regulators of the *BRC1* network (TF2Network tool^25^) also predicted some of these BTFs as potential regulators of the network (i.e. ABI5, GBF2, GBF3, ABF3, ATAF1, MYBD, HB21, HB40, HB53; **Supplemental Dataset 5**).

To investigate whether the BTFs targeted the *BRC1*-dependent genes, we studied their genome-wide binding sites using publicly available ChIP-seq and DAP-seq data^17,24^. Remarkably, BTF targets were significantly enriched in *BRC1*-dependent BRC1-bound, and *bona fide* BRC1 targets (**Fig. 5a**). Fine mapping of binding peaks revealed that in genes bound both by BRC1 and a BTF, a large proportion of the BRC1 peaks co-occur with BTF peaks (**Fig. 5b**). This suggests that BTFs act in close cooperation with BRC1 to control transcription. Furthermore, not one but several BTF peaks usually overlap with BRC1 peaks at the promoters of BRC1 *bona fide* gene targets (**Fig. 6b,c**). In the *BRC1*-dependent gene targets not bound by BRC1, the peaks of the BTFs also overlap (**Fig. 6d)**.

**Fig. 6.**
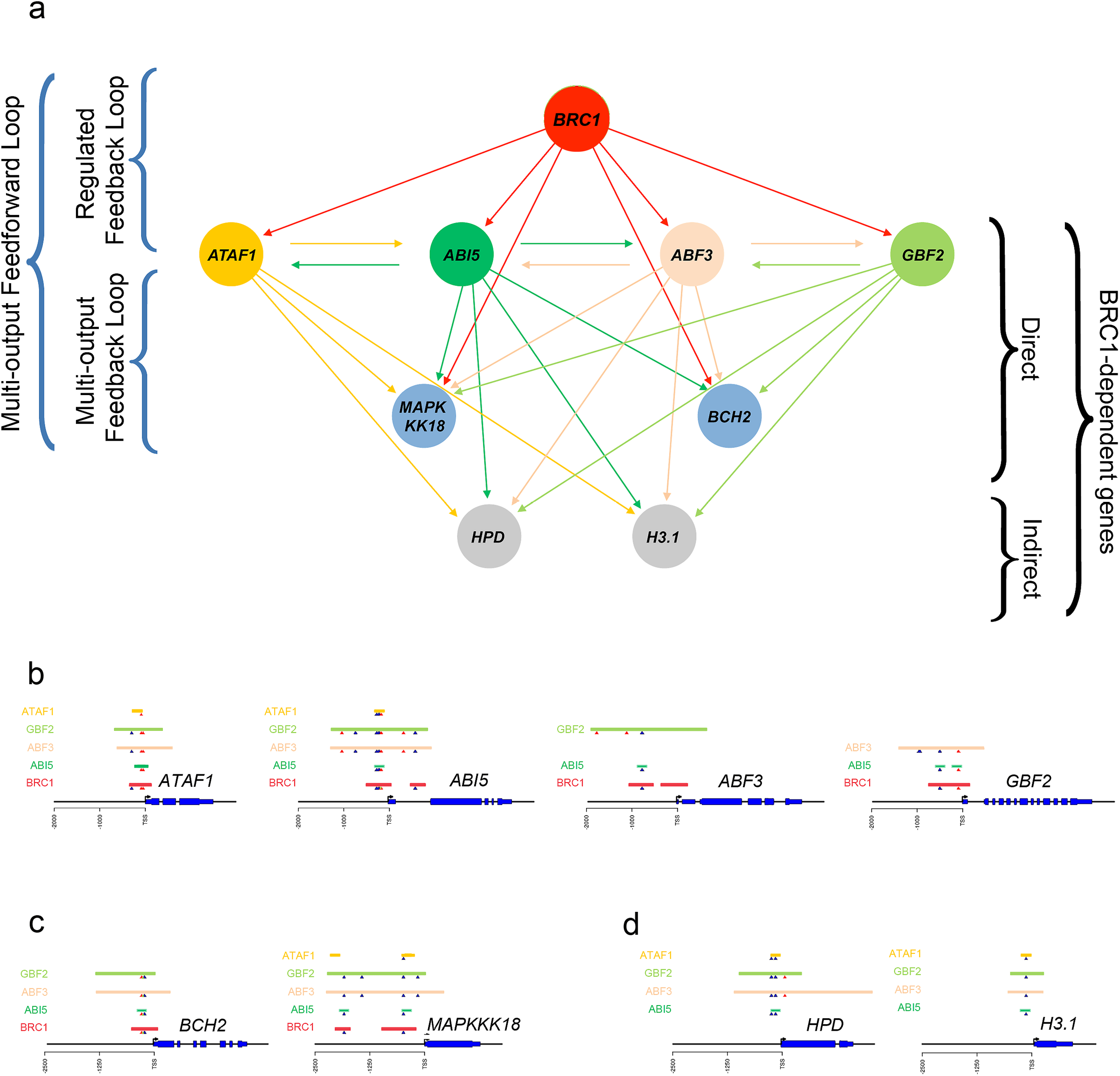
Motifs overrepresented in the BRC1 network. **a**, Simplified representation of the BRC1 network, that exemplifies overrepresented motifs (multi-output FFLs, regulated FBLs and multi-ouput FBLs) using specific cases. Orange and green circles represent genes encoding BTFs (ATAF1, ABI5, ABF3, GBF2); blue circles, other *bona fide* BRC1 targets; grey circles, *BRC1*-dependent genes bound by BTFs but not by BRC1. Green and orange arrows indicate direct binding of the BTFs; red arrows, direct binding of BRC1. **b**, Genomic region of genes encoding BTFs. The overlapping binding sites of BRC1 and BTFs are represented. **c**, Genomic regions of *bona fide* BRC1 direct targets *BCH2* and *MAKKK18*; in their promoters BRC1 and BTFs binding sites overlap. **d**, Genomic region of *BRC1*-dependent genes *HDP* and *H3.1*. Their promoters are not bound by BRC1 but they are bound by all four BTFs, whose binding sites overlap.

### The BRC1 transcriptional network is robust against mutation and enriched in multi-output feed-forward and regulated feed-back loops

To further explore the regulatory interactions between BRC1, BTFs and other TFs, we constructed and analysed a transcriptional network integrating our BRC1 ChIP-seq data with ChIP-and DAP-seq data publicly available for ten BTFs and 27 additional *BRC1*-dependent TFs^17,24^. The network comprises the 947 *BRC1*-dependent genes (**Supplemental Dataset 1**). This transcriptional network, BRC1NET, can be accessed and analysed using an interactive web application available at https://greennetwork.us.es/BRC1NET/. The BRC1NET application allows dynamic visualization of specific and common targets for any TF or combination of TFs in the network. It generates target gene lists, performs significance and functional enrichment analyses, and allows visualization of TFs binding peaks and binding motifs in any given gene of the network (**Supplemental Fig. 6**).

Using the BRC1NET enrichment analysis tool, we found that the targets of some BTFs are significantly enriched in genes of particular *BRC1*-dependent clusters, which suggests that they can largely contribute to the expression of specific *BRC1*-dependent clusters (**Supplemental Tables 2,3, Supplemental Fig. 7**). For instance, GBF2, GBF3, ABF3 and ANAC032 could mainly regulate the ethylene-related cluster UP_C1; ABF3, ABI5 and ANAC032 could regulate the ABA-related cluster UP_C6. The DOWN clusters were not significantly targeted by any BTF.

Next, we performed a network motif enrichment analysis of BRC1NET, to identify non-random *gene modules* formed by TFs that co-ordinately orchestrate the regulation of common target gene sets (**Supplemental Dataset 6**). We found that this network is enriched in multi-output feed-forward loops (FFLs), and regulated multi-output feedback loops (FBLs) (**Supplemental Fig. 8**)^26^.

In multi-output FFLs, a master regulator TF1 binds the promoter of a gene encoding and intermediary regulator TF2. Both TFs bind the promoters of a common target gene set, thus regulating their expression (**Supplemental Fig. 8d**). These network motifs are found in transcriptional networks in which environmental signals induce rapid and robust, but reversible, responses^27,28^. In BRC1NET, a large number of FFLs comprise BRC1 as a master TF1, and a BTF (i.e. ABI5, ANAC032, ATAF1, HB21, HB40, HB53, GBF2) as TF2. BRC1 and the BTF in turn regulate common target gene sets (**Fig. 6a, b**).

The BTFs are also involved in BRC1-regulated FBLs with multiple outputs: in these network motifs two BTFs (TF1 and TF2), regulate each other, and jointly control a set of common target genes (**Fig. 6a-d, Supplemental Fig. 8b,c**). These motifs are commonly found in transcriptional networks controlling developmental cell fates, and can transform a transient signal into a stable response. The targets of these FBLs are not randomly distributed over the network. Rather, pairs of BTFs also involved in FBLs, significantly regulate specific clusters (**Supplemental Tables 3,4)**. For instance, the ABI5/ANAC032 FBL target gene set is significantly enriched (p-val 5.7×10^−12^) in genes of the ABA-related cluster UP_C6 (**Supplemental Fig. 9**), whereas the ABI5/ATAF1 FBL target gene set regulates cluster UP_C2 (p-val 5.3×10^−5^). Interestingly, no FFLs or FBLs were identified that significantly regulate the DOWN clusters, which supports the position of these clusters downstream of but far away from the direct signalling response triggered by BRC1.

This intricate *BRC1*-dependent network is highly redundant: a significant proportion of the 947 *BRC1*-dependent genes are targets of more than one BTF (e.g. **Fig. 6b, c**), which renders the network extremely robust to random mutations.

Indeed, the network connectivity is only mildly affected even after *in silico* removal of up to 10 TFs of the network (excluding BRC1, **Supplemental Fig. 10)**, which implies that a plant carrying mutations in all 10 of these TFs will likely still have a wild-type branching phenotype under standard conditions.

All these results suggest that BRC1 regulates a transcriptional network robust against noise and mutation, reversible in the first steps consisting of FFLs, but stable once stablished by FBLs. BRC1 controls this network both directly, by forming FFLs and regulated FBLs with the BTFs (**Fig. 6a,b,c**), and indirectly, via numerous FBLs established among the BTFs and other *BRC1*-dependent genes not bound by BRC1 (**Fig. 6a,d**).

## Discussion

We show here that *BRC1* is a master regulator of a transcriptional network that promotes growth inhibition in axillary buds. BRC1 directly controls the expression of a collection of BTFs that modulate and amplify *BRC1* function, both by binding other BRC1 direct targets, and by regulating secondary *BRC1*-dependent genes not bound by BRC1.

One of the main mechanisms by which *BRC1* promotes dormancy is by induction of ABA signalling to cause growth arrest. We had previously shown that BRC1 directly regulates the expression of *HB21, HB40* and *HB53*, with in turn activate (together with *BRC1) NCED3*, an enzyme catalyzing a rate-limiting step of ABA biosynthesis^6^. Now we demonstrate that BRC1 has a more pervasive influence on ABA signalling: a significant proportion of the *bona fide* BRC1 direct targets are involved in ABA metabolism, transport, perception, signalling and response (**Supplemental Dataset 4)**. This enrichment in ABA-related targets has also been observed for maize TB1^7^. Indeed, several common targets between TB1 and BRC1 are related to ABA (e.g. *HB53/HB40, ANAC032, ABF3, ABI5, NCED3, LEA4-5*, **Supplemental Results**). This suggests that the control of ABA signalling by BRC1/TB1 genes is evolutionary ancient and predates the separation of monocot and dicotyledoneous plants. BRC1 may also induce GA degradation (via induction of *GA2OX2*), which would stabilise the growth-inhibiting DELLA proteins. Furthermore, BRC1 directly upregulates the SL biosynthesis gene *MAX1*. SL signalling boosts *BRC1* mRNA accumulation thus probably creating a positive regulatory feedback loop that bolsters *BRC1* responses and bud dormancy.

Our ChIP-seq and RNA-seq data show that BRC1 mainly acts as a transcriptional activator, which binds GDCCCA DNA motifs preferentially located in gene promoters, in close proximity to the TSS. Likewise, TB1 binds a related (although not identical) motif (GGnCCC) mainly in gene promoters^7^. However, the significant enrichment of BRC1 peaks in G box-like motifs (BACGTGKC), suggests that BRC1 targets are co-regulated by G box-binding factors. Notably, ABA transcriptional networks are also enriched in G box-containing <u>AB</u>A <u>R</u>esponse <u>E</u>lements (ABREs)^29^, and several BTFs (i.e. bZIP factors ABI5, GBF2, GBF3, ABF3, NAC factors ANAC032 and ATAF1) bind G-box motifs and are ABA master regulators^24,29,30^. Furthermore, the specific gene targets and binding peaks of the BTFs significantly overlap with those of BRC1, which makes them likely candidates to have a critical impact on the *BRC1* transcriptional network. BRC1 and G-box binding motifs are in some cases in close proximity, raising the possibility that BRC1 and BTFs physically interact. High-throughput CrYTH-seq assays revealed a potential interaction between BRC1 and ABI5^31^ but this has not yet been confirmed *in vivo*. Notably, although *BRC1* expression is sufficient to activate BTFs and their downstream targets^6^ (this work), these genes also belong to a well-characterised ABA network, induced in tissues where *BRC1* is not usually expressed (e.g. ABA-treated seedlings^24,32^. This strongly suggests that BRC1 has co-opted a pre-existing ABA signalling network to be activated (under BRC1 control) in axillary buds. This could have been achieved by acquisition of BRC1 binding sites at promoters of top-tier ABA master regulators to become BTFs (i.e. *ABI5, ABF3, GBF2, GBF3, ANAC032, HB21, HB40, HB53*), which could in turn activate additional ABA-related targets, including other TFs. BRC1 binding sites also evolved in these secondary targets creating FFLs between BRC1 and the BTFs. The observation that BRC1 is not induced by ABA treatments^6^ and is not co-expressed with ABA networks in other tissues, supports the view that BRC1 is an ABA-independent, master regulator of the ABA network, exclusively in buds. Perhaps BRC1 increases chromatin accessibility of the BTFs to their targets in axillary buds, a tissue in which otherwise BTFs might not bind their targets. Some BTFs (e.g. ABF3, ABI5, ATAF1) require ABA-induced phosphorylation for full activation^30^. *BRC1-*mediated ABA biosynthesis (by upregulation of *NCED3* and *BCH2*) may be instrumental for this post-translational activation, and for additional support of the ABA-related responses.

The BRC1 transcriptional network is enriched in multi-output FFLs^26^ formed by BRC1, BTFs and *bona fide* BRC1 targets. FFLs regulate reversible decisions and act as sign-sensitive delay elements and persistence detectors. Some target genes may require both BRC1 and a BTF for induction. These would show delayed responses to *BRC1*, but no delay when *BRC1* is downregulated. This would help filtering out signals that transiently induce *BRC1* expression, such as a brief shading. Only persistent signals (e.g. a dense plant canopy) would lead to network induction. In contrast, other genes could be induced by either BRC1 or a BTF. These targets would have a rapid and persistent response even during a transient loss of *BRC1* induction. BRC1NET is also enriched in multi-output FBLs formed by the BTFs - which regulate each other- and jointly control downstream genes. Unlike FFLs, FBLs provide memory of an input signal even after the signal is gone. They are frequent in transcriptional networks that transduce signals into stable developmental decisions. In BRC1NET, these motifs could ensure that environmental or endogenous signals upregulating *BRC1*, can then be locked into a steady state-response that stabilises bud dormancy. The final output will depend on the prevailing network motifs, their sign (positive or negative), and additional, *BRC1*-independent regulation of network components. It is plausible that the degree of reversibility of bud dormancy (i.e. para- and ecodormancy vs endodormancy) is in part determined by the prevalence of FFLs that allow delayed responses and reversibility, or FBLs that promote more stable and lasting responses of growth arrest. The BRC1NET is very robust against mutation, an essential feature to maintain a high fitness state. In line with this, the network connectivity is only mildly affected even after removal of up to 10 TFs (**Supplemental Fig. 10)**.

In summary, during the evolution of angiosperms, BRC1/TB1 genes seem to have recruited an ABA-related network comprising a collection of master regulators of ABA signalling and their targets. This highly connected network, robust against noise and mutation, constitutes a transcriptional cascade that in the first steps is reversible and filters out transient signals (using FFLs), but becomes stable once established (due to the FBLs). The flexibility of this network is essential for an adaptive response and a correct carbon allocation in the plant under growth-limiting conditions. This knowledge will prove fundamental to optimise plant architecture and crop production.

## Materials and methods

### Plant materials and growth condition

Wild-type Arabidopsis thaliana plants were of the Columbia-0 (Col-0) ecotype. The brc1-2 mutant was described^10^ and the *GFP:BRC1^ind^* lines have been generated previously^6^. All lines were grown on 1/2MS medium 1% (w/v) agar plates under long day conditions (16/8 light/dark cycle) at 21°C.

### Promoter:GUS constructs

Genomic fragments of 1728, 1709, 1830, 2265, 1846, 540, and 1990 base pairs comprising the BRC1 binding peaks upstream the ATG of *ABI5, ABF3, GBF2, GBF3, MYB3, ATAF1* and *NAP*, respectively, were amplified from genomic DNA using primers in **Supplemental Table 5** and Taq Phusion polymerase (New England BioLabs), cloned in pGEM-t Easy (Promega), re-amplified by PCR with primers with attB tails and BP-cloned into pDONR207 (Invitrogen). Promoter:GUS binary vectors were generated by LR recombination using Gateway LR clonase II (Invitrogen) according to manufacturer’s instructions into the destination vector pGWB3.

### Generation of transgenic plants

Binary vectors were transformed into the Agrobacterium tumefaciens strain AGL-0. Transgenic Arabidopsis plants were generated by agroinfiltration using the floral dip method^33^. At least 10 T3 homozygous lines were generated and analysed for each construct.

### GUS staining

GUS staining was conducted as described in^34^.

### RNA-seq and ChIP-seq material collection

For the RNA-seq experiment, plants of *brc1-2* (control) and *GFP:BRC1^ind^*lines were grown on plate. Ten-day old seedlings were induced with 5mL of 10μM estradiol per plate. After an induction period of five hours, 1 g of tissue was harvested for each of the three biological replicates. Total RNA was isolated using the InviTrap^®^ Spin Plant RNA Mini Kit according to the manufacturer’s protocol. TURBO™ DNase was used to clean the RNA samples from DNA. Library preparation for whole genome RNA sequencing was done using the Illumina Truseq Library Preparation Kit. Library quality was evaluated using a Bioanalyzer and an RNA Nano 6000 kit (Agilent). RNA concentrations were determined using the Xpose ‘DSCVRY’ (Trinean). The libraries were then sequenced on the Illumina Hi-Seq 2500.

For the ChIP-seq experiment, plants of *GFP:BRC1^ind^* lines were grown on plate. Ten-day old seedlings were induced with 5mL of 10μM estradiol per plate. After an induction period of five hours, 1.5g of tissue was collected per sample. ChIP was performed as described^35^, using μMACS Anti-GFP beads (Miltenyi). Input DNA was used as control, which is isolated from the sonicated chromatin prior to immunoprecipitation.

### Identification of Genes Coregulated with *BRC1*-dependent genes

BRC1-dependent genes were obtained from^4^ Coregulation of the 181 UP and 57 DOWN was analysed by hierarchical clustering based on 15,200 publicly available microarray experiments (Hcluster, ATTED-II^36^). Additional coregulated genes were obtained with CoEx-Search (ATTED-II^36^). The expression of these genes was validated in the original arrays ^4^. Only genes upregulated (positive fold change FC) in WL+FR treated wild type dormant buds but not in WL treated or brc1 mutant buds were included in the extended list of *BRC1*-dependent genes.

### Functional Annotation of *BRC1*-dependent clusters

GO automated analyses of function prediction were carried out using a statistical overrepresentation test (pval < 0.05) followed by Bonferroni correction for multiple testing in the PANTHER classification system^8^.

### Differential gene expression and gene ontology (GO) enrichment analyses

Libraries of three biological replicates were sequenced individually and analysed using the Bowtie–Tophat–Cuffdiff (BTC) pipeline^37^. Differential gene expression was determined for all estradiol induced *GFP:BRC1^ind^* samples, using induced *brc1-2* samples as control. The cut-off was set at a false-discovery rate (FDR) <0.05 in all analyses performed. The RNA-seq data is made available via NCBI and can be accessed through GEO accession number GSE155028. The BINGO 3.03 plug-in^38^, implemented in Cytoscape 2.81^39^, was used to determine and visualise the Gene Ontology (GO)-enrichment categorization. A hypergeometric distribution statistical testing method was applied to determine the enriched genes in combination with the Benjamini and Hochberg multiple testing correction (FDR<0.05).

### ChIP-seq data analysis

Libraries of three biological replicates of both the ChIP and Input samples were sequenced after which Bowtie2 was used to map the reads to the Arabidopsis genome ^40^. MACS2 was used for peak calling (the statistical detection of protein binding sites in the DNA)^41^ using the Input samples as control. The ChIP-seq data is made available via NCBI and can be accessed through GEO accession number GSE155028. The function Distance2Genes, part of the R-package ‘CSAR’, was used to determine the nearest gene^42^. These nearest genes are then presented in a list of candidate target genes^35^ for further analysis. The analysis of read distribution of our ChIP-seq Dataset was done by the R-package ChIPpeakAnno: ‘assignChromosomeRegion’. As a cutoff 1kb of promoter region with an immediate cutoff of 0.5 kb was used. All three biological replicates were initially analysed separately. To confirm the high rate of reproducibility the overlap between the two highest enriched samples and the overlap of all three samples was analysed. To assess the reproducibility of the datasets, a pair-wise correlation analysis was done, revealing Pearson correlation co-efficiencies ranging from 0,77 to 0,83 (**Supplemental Fig. 3b**) which supports the reproducibility and quality of our experiments^44^. Because of the lower number of peaks in sample 3 (**Supplemental Fig. 3a**), we decided to perform further analyses on the first two ChIP-seq datasets. We approached these datasets by creating a file representing their overlap, which was inferred by the distance between peaks in the individual samples. The average peak size was ±300bp, so if the centre of a peak in one sample is no more than 150bp apart from the centre of a peak in the other sample, the peaks are assumed to be in fact the same and were saved in a separate file for subsequent analyses. This criterion resulted in a file containing roughly 60-70% of the peaks of sample 1 and 2. Since this merging of peaks may result in a slight shift of the actual binding site, analyses looking specifically at the exact location of the binding site were done on the individual datasets.

### Analysis of peak-overlap between BRC1 and BTFs

We use as datasets the 4227 BRC1 peak locations and the peak locations and target genes for 10 BTFs. We defined promoter regions as 3kb upstream region according to TAIR. For each BTFs we performed the following analysis: we obtained the promoter region of the BTF target genes; retrieve the peak locations of BTF occurring in those promoter regions (at least one bp of the peak should be in the promoter region); obtain the peak locations of BRC1 occurring in those promoter regions; for each BRC1 peak occurring in those promoter regions, check if there is a BTF overlapping peak (defined as having at least one bp in common).

### Motif enrichment analysis

Motif discovery was carried out using MEME-ChIP^45^. This program consists of several sub-programs that each perform a specific analysis. In the present study MEME, DREME and CentriMO were used. MEME^46^ looks for overrepresented motifs in a set of sequences compared to a background model of nucleotide frequencies. The background model was generated using fasta-get-markov for the MEME suite, using a second order Markov model, and a set of 207 randomly selected sequences from Arabidopsis promoters as input. DREME^15^ looks for motifs occurring significantly more often in the input set of sequences compared to a control set; as control set, shuffled versions of the input set was used. CentriMO^47^ is used to test if the motifs found are centrally enriched. In the present study, motifs were investigated if they were in the top six motifs as defined by MEME-ChIP. Defaults settings for DREME were used, including an E-value threshold of 0.05. Default settings for MEME were used, with the following exceptions: the setting meme-mod-anr was used, which assumes zero, one or multiple motif occurrences per sequence. Minimum and maximum width for the motifs were set to 4- and 15bp, respectively.

### Network construction and visualization

The transcriptional network comprising the *BRC1*-dependent genes and the potential targets of the BRC1-dependent TFs was constructed using the R package igraph for network construction and analysis and ChIP-seq and DAP-seq data available^17,24^. Nodes represent genes, and edges are drawn from gene_i_, to gene_j_ when gene, encodes a TF that binds to the promoter of genej. A network adjacency matrix of order 947×947; number of rows (genes) x number of columns (genes); was constructed by setting component (i,j) to 1 when gene_i_, encodes a TF that binds to genej promoter. Otherwise it was set to 0. The function graph.adjacency with input the previously described adjacency matrix and mode equal to directed was used to generate our network comprising 947 nodes/genes and 10240 edges/potential direct transcriptional regulations. A web app was developed using the R package shiny to analyse this network (https://greennetwork.us.es/BRC1NET/)

### Network motifs identification and analysis

A network motif is defined as a subgraph whose occurrence is significantly greater in our network than its occurrence in random networks with the same topological properties. The analysis was performed using the R package igraph. 1000 random networks were generated with the same number of nodes as our transcriptional network, namely 947 nodes. To capture the same topology as our network, 37 nodes were chosen randomly that would play the role of the TFs in our network. Each of these nodes was connected to the same number of nodes as one of the TFs in our network, but the targets were also chosen randomly.

## Supporting information

Supplemental Results and Figures

## Supplemental Information

Supplemental Information includes Supplemental Results, Supplemental figures and Supplemental tables.

## Acknowledgements

The research was supported by grants from the Dutch Scientific Organization (NWO); (NWO-JSTP grant 833.13.008 and VENI 15060), Spanish Ministry of Economy (MINECO) and fondos FEDER [grants BIO2014-57011-R, BIO2017-84363-R and BIO2017-84066-R]. CT was a La Caixa fellow and Aitor Muñoz a FPI (MINECO) fellow. We thank Juan Carlos Oliveros, Mónica Franch and F.J.M. van Workum for help with the RNA- and ChIP-seq data analysis, Estíbaliz Bustos for amplification of BTF promoters and Javier Paz-Ares and Desmond Bradley for constructive criticisms of the manuscript.

