## Supplemental Results and Figures for "A gene regulatory network critical for axillary bud dormancy directly controlled by Arabidopsis BRANCHED1"

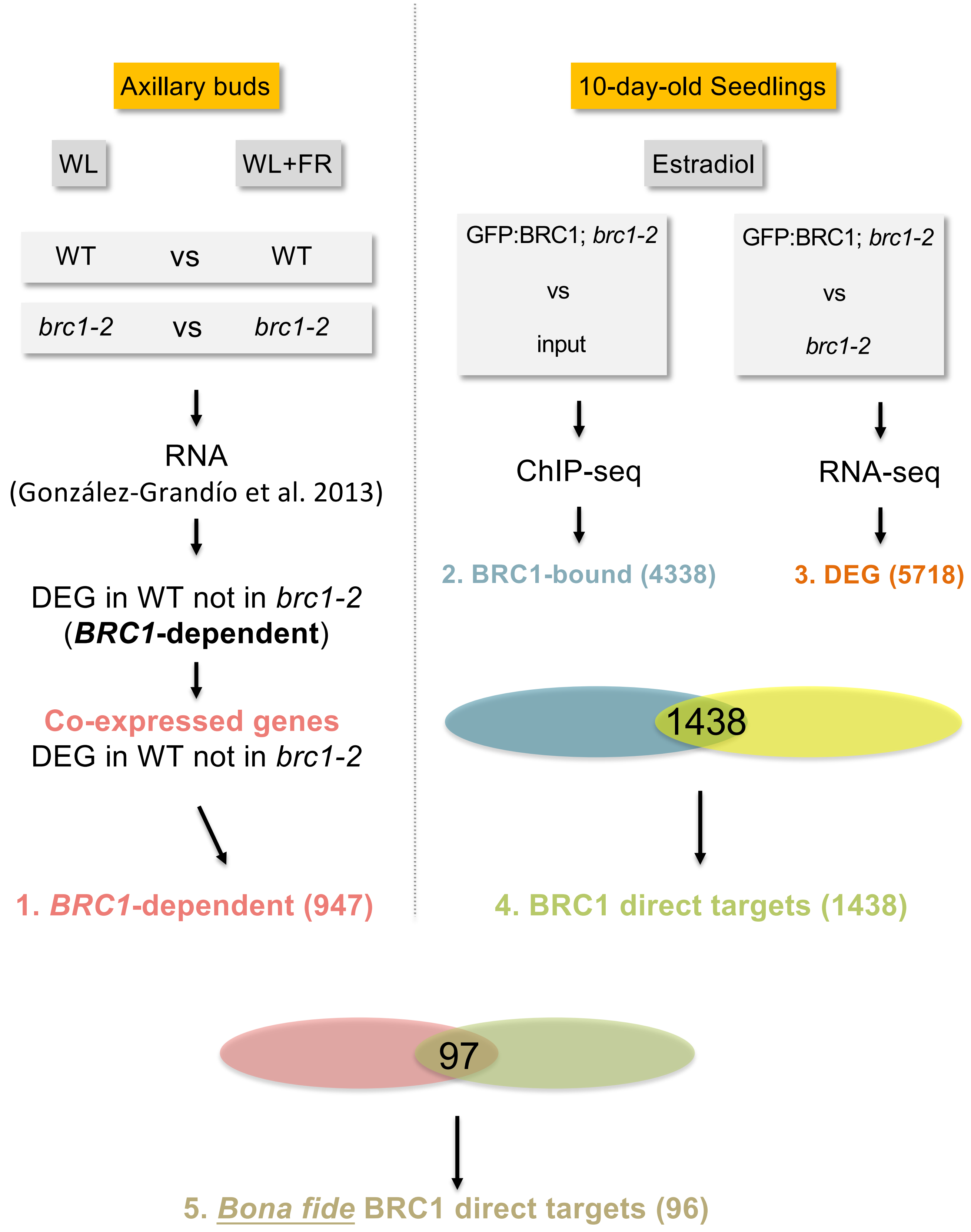


**Supplemental Fig. 1.** Schematic representation of the steps followed to identify *bona fide* BRC1 direct targets. Terms used in this paper (i.e. *BRC1*-dependent, BRC1-bound, BRC1 direct targets, and *bona fide* BRC1 direct targets) are indicated.

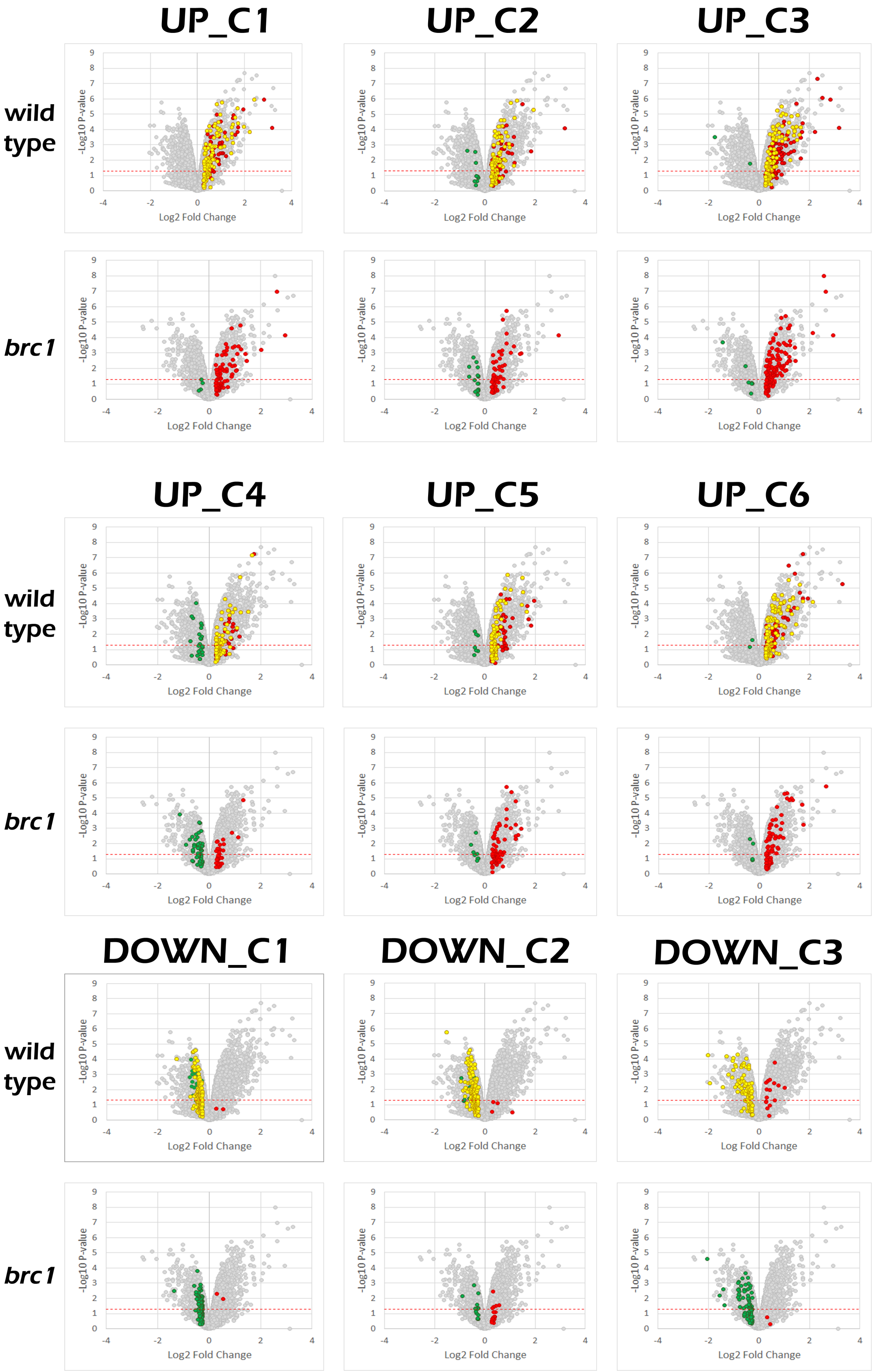


**Supplemental Fig. 2** Volcano plots representing representing p-val (–Log10 pval, vertical axis) and relative expression (Log2 Fold Change, horizontal axis) of genes in the active vs dormant buds (8 h low R:FR vs. High R:FR) microarray. Yellow, *BRC1*-dependent genes (DEG in wild type but not in *brc1* mutants). Red, genes upregulated both in wild type and *brc1* mutants. Green, genes downregulated both in wild type and *brc1* mutants. Only *BRC1*-dependent genes (yellow) were used for subsequent analyses.

A)

| | Sample | Illimina Hiseq 2000 Total # reads | Quality Check 35N | % 35N | Bowtie2 read alignment # reads mapped | % total | Sample | MACS2 peak calling # peaks | Overlap 1-2 | Overlap 1-2 | Overlap 1-2-3 | Overlap 1-2-3 |
|---|---|---|---|---|---|---|---|---|---|---|---|---|
| ChIP | 1 | 32.270.369 | N/A | N/A | 7.163.351 | 22,2 | 1 | 4567 | 2761 | 60% | | 8% |
| | 2 | 34.830.313 | N/A | N/A | 11.764.490 | 33,78 | 2 | 3951 | | 70% | 374 | 9% |
| | 3 | 32.332.938 | N/A | N/A | 2.988.264 | 9,24 | 3 | 450 | | | | 83% |
| Input | 1 | 13.544.249 | N/A | N/A | 13.405.792 | 98,98 | | | | | | |
| | 2 | 17.273.742 | 18.988 | 0,11 | 17.126.011 | 99,14 | | | | | | |
| | 3 | 3.055.942 | 3.634 | 0,12 | 3.004.333 | 98,31 | | | | | | |

B)

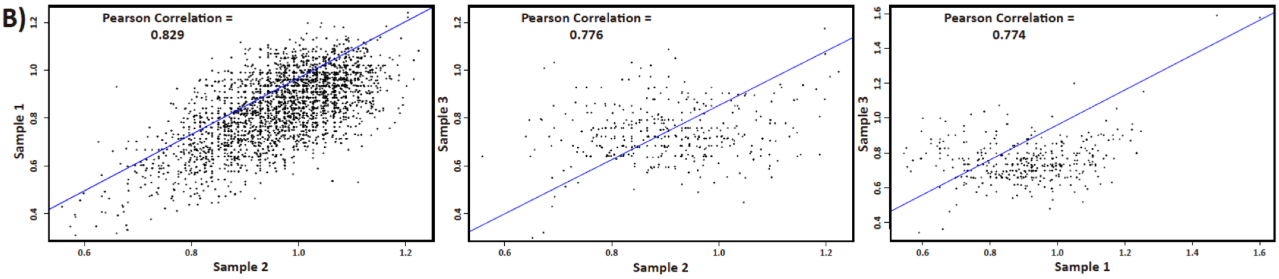


**Supplemental Fig. 3 Analysis of BRC1 ChIP-seq datasets. a,** Summary of the sequencing results. From left to right: total number of reads; number and percentage of failed reads (35N, %35N); number and percentage of reads mapping to the Arabidopsis genome; number of peaks significantly enriched; number and percentage of overlapping peaks between samples 1 and 2; number and percentage of overlapping peaks between samples 1, 2 and 3. **b**, Pearson Correlation Coefficients of plots comparing all three biological replicates.

| | Sample 1 | | | | Sample 2 | |
|---|---|---|---|---|---|---|
| | **Motif** | **E-value** | | | **Motif** | **E-value** |
| 1. | BACGTGKC | 4.1e-403 | | 1. | BACGTGKC | 3.5e-337 |
| 2. | GDCCCA | 6.1e-212 | | 2. | GDCCCA | 1.0e-185 |
| 3. | AGAGAGAR | 2.0e-149 | | 3. | AAAANARA | 1.6e-130 |
| 4. | CACRYG | 2.9e-122 | | 4. | BAGAGAS | 2.1e-106 |
| 5. | DAAGAARA | 8.4e-122 | | 5. | CACRYG | 5.8e-101 |
| 6. | AAAHAAAA | 1.7e-077 | | 6. | AGAAGANR | 9.6e-082 |
| 7. | AGAGAVR | 4.3e-073 | | 7. | GTGGNS | 8.9e-058 |
| 8. | GTGGNS | 3.0e-065 | | 8. | BTATATA | 3.2e-052 |
| 9. | TATATAB | 1.3e-058 | | 9. | CTCTYY | 6.2e-032 |
| 10. | TATTATW | 1.3e-030 | | 10. | AATAMTAW | 2.0e-027 |
| 11. | RACMAAA | 8.5e-023 | | 11. | ATATAT | 5.0e-018 |
| 12. | ACGTGK | 3.5e-022 | | 12. | GGDCCA | 2.2e-017 |
| 13. | AGCCCAW | 2.7e-021 | | 13. | MCACACA | 1.1e-015 |
| 14. | AYAMATA | 6.7e-020 | | 14. | AAASMAAA | 8.1e-014 |
| 15. | CTCTYY | 8.0e-018 | | 15. | ACGTGK | 1.2e-013 |
| 16. | MCAAGTGK | 6.7e-017 | | 16. | ATARAKA | 1.9e-013 |
| 17. | GGDCCA | 5.3e-016 | | 17. | MCAAGTGK | 1.1e-012 |
| 18. | CKTCTYC | 9.6e-013 | | 18. | CGACGNCG | 1.2e-011 |
| 19. | AGCYGTC | 3.7e-012 | | 19. | AGCCCAW | 8.4e-010 |
| 20. | AAAGAAAS | 2.3e-008 | | 20. | GAAAANGA | 1.1e-008 |
| 21. | CACWCACA | 1.5e-008 | | 21. | RAGAAGS | 3.5e-008 |
| 22. | CRTCATCA | 1.0e-007 | | 22. | GCCGHC | 2.4e-006 |
| 23. | ADATAT | 3.1e-006 | | 23. | AGCCGTTR | 1.2e-005 |
| 24. | GAWAAMGA | 6.0e-007 | | 24. | TACTASTA | 1.4e-005 |
| 25. | CAACGGCK | 7.7e-006 | | 25. | AGCTGKC | 4.5e-005 |
| 26. | CGHCGAC | 1.9e-005 | | 26. | TACRTAYA | 3.8e-005 |
| 27. | TACTASTA | 8.8e-005 | | 27. | AAAGHCAA | 5.1e-005 |
| 28. | TAATTAR | 9.8e-005 | | 28. | AGAGA | 3.9e-005 |
| 29. | CAAAVCC | 1.1e-004 | | 29. | GATGAYGA | 3.7e-004 |
| 30. | RGACAAGA | 5.8e-004 | | 30. | AAAATATV | 5.8e-004 |
| 31. | CMACTTGC | 1.1e-003 | | 31. | BCTCCTCC | 7.0e-004 |
| 32. | CGGCGGM | 1.9e-003 | | 32. | CCGACCCG | 1.5e-003 |
| 33. | GGGTCGGK | 2.8e-003 | | 33. | GGGACM | 2.3e-003 |
| 34. | AAGCHAA | 8.3e-003 | | 34. | CCACYA | 2.6e-003 |
| 35. | GAGAGA | 1.1e-002 | | 35. | CGSCGTTA | 8.3e-003 |
| 36. | CAAAGACR | 3.6e-002 | | 36. | AACRACAA | 2.3e-002 |
| 37. | GGAGGAGR | 3.9e-002 | | 37. | ATTATTGD | 4.9e-002 |

**Supplemental Table 1.** Overview of significantly overrepresented motifs in BRC1 ChIP-seq peaks. Motifs have been determined by the DREME (Discriminative Regular Expression Motif Elicudation) script, which is part of the MEME Suite (Motif Sequence Analysis Tools).

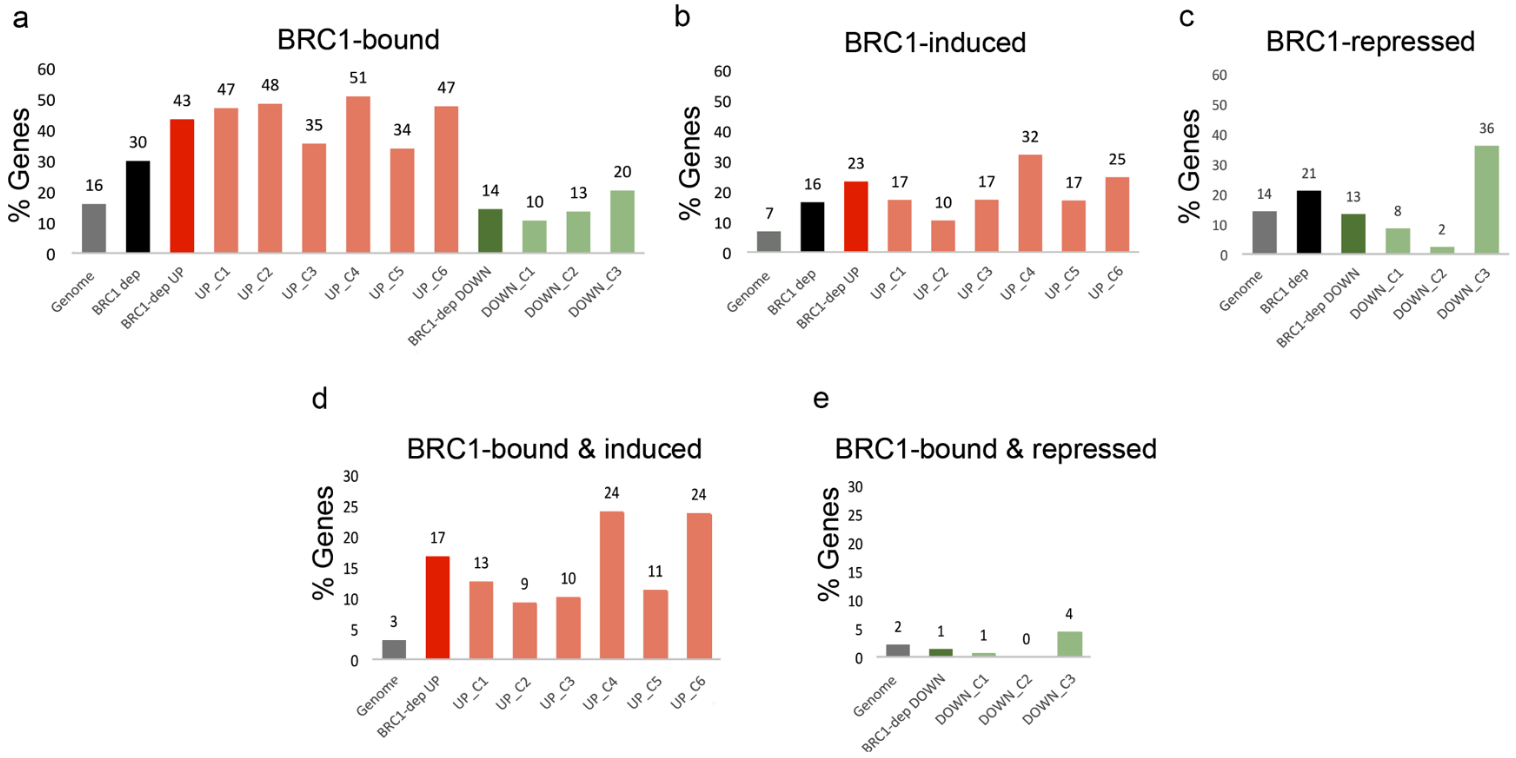


**Supplemental Fig. 4. The UP *BRC1*-dependent clusters are enriched in BRC1 direct targets.**
**a**, Percentage of BRC1-bound genes in the Arabidopsis genome (16%) and in each of the BRC1-dependent clusters. **b**, Percentage of induced genes in the Arabidopsis genome (7%) and in each of the BRC1-dependent clusters. **c**, Same as **b**, but for repressed genes. Percentage of BRC1 direct targets (BRC1-bound & differentially expressed) in the *BRC1*-dependent UP (d) and DOWN (e) clusters.

| TF | Motifs | Motif E-value | Motifs | Motif E-value |
|---|---|---|---|---|
| GBF2 | 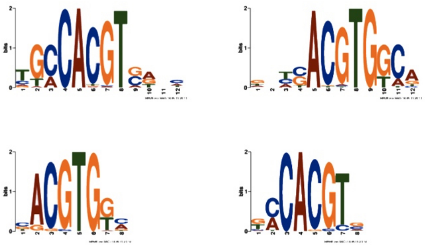 | 0<br>1.1e-246 | 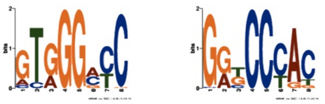 | 1.9e-15 |
| GBF3 | 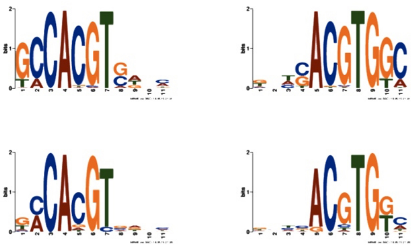 | 0<br>7e-221 | 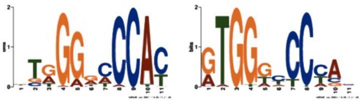 | 3.7e-22 |
| ABF3 | 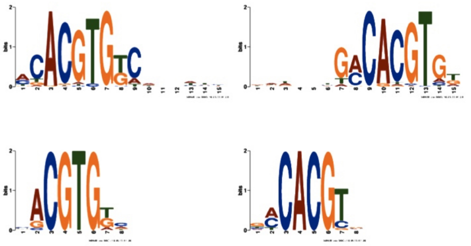 | 0<br>4.4e-82 | | |
| ANAC032 | 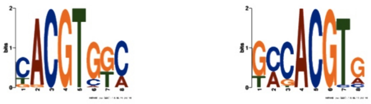 | 7.3e-07 | | |
| ABI5 | 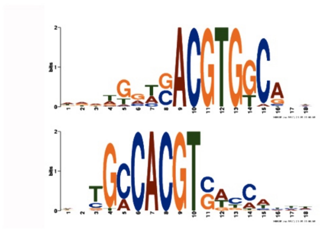 | | | |
| ATAF1 | 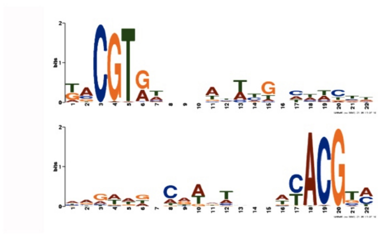 | | | |

**Supplemental Fig. 5. Motifs enriched in the binding sites of BTFs indicated**. Left column, G-box like motifs. Right Column, TCP-like motifs. GBF2, GBF3, ABF3 and ANAC032 motifs were obtained from ChIP-seq experiments carried out in seedlings treated with 10 mM ABA for 4 h (Song et al. 2016). ABI5 and ATAF1 data motifs were obtained by DAP-seq data (O´Malley et al. 2016).

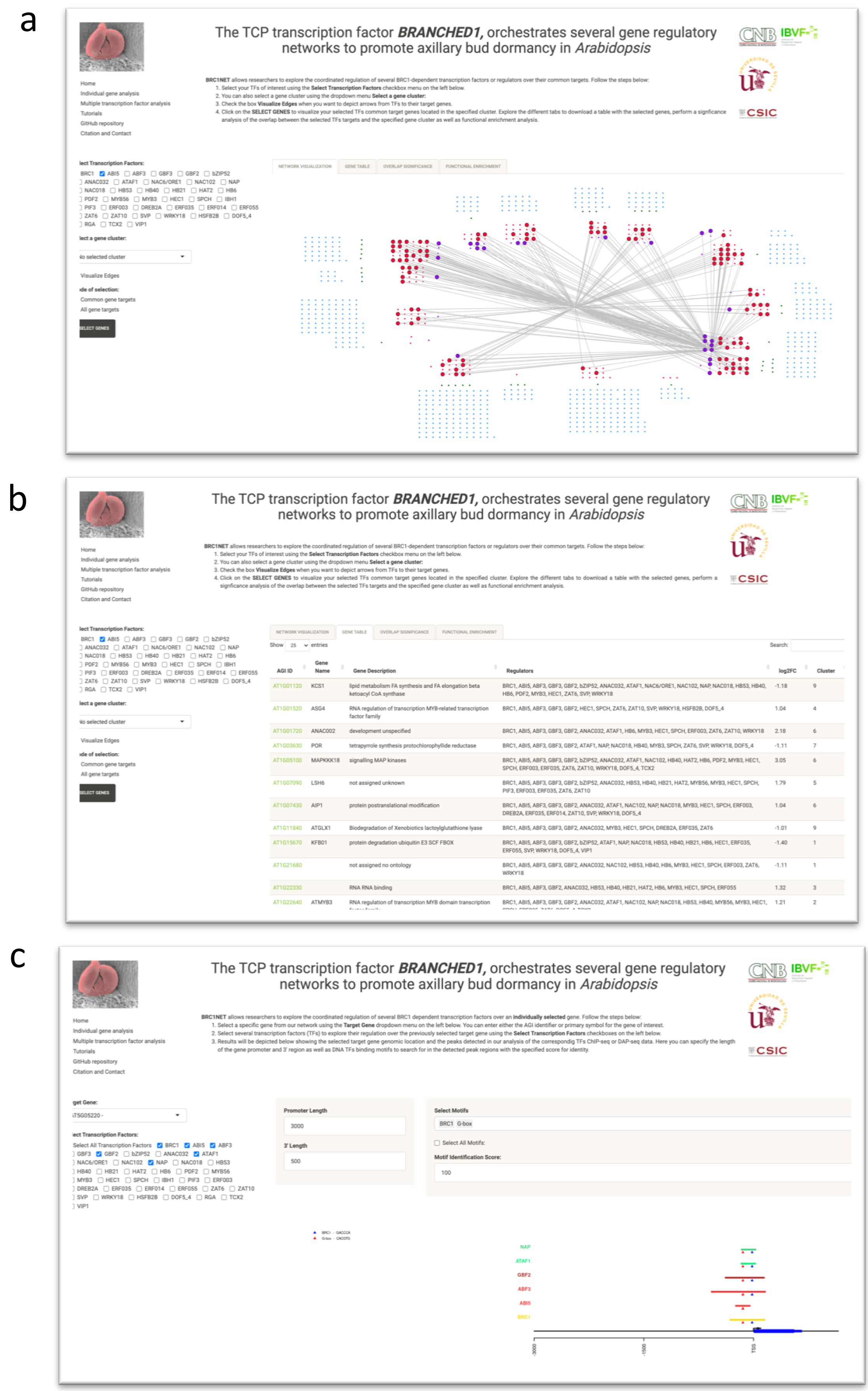


**Supplemental Fig. 6. Screenshots of the BRC1NET web application.** **a**, In this example the Multiple Transcription Factor Analysis tool is used to search for BRC1 and ABI5 common targets. Dots represent network nodes, grouped by *BRC1*-dependent clusters. Edges link BRC1 and ABI5 with their direct targets. **b**, Multiple Transcription Factor Analysis tool, displaying a list of the target genes selected. **c**, Individual Gene Analysis tool displaying binding sites of selected TFs at the genomic region of one *BRC1*-dependent gene. 5´and 3´genomic lengths can be adjusted. DNA binding motifs for all TFs available can also be displayed. BRC1NET is available at https://greennetwork.us.es/BRC1NET/

| BTF/Cluster | UP_C1 | UP_C2 | UP_C3 | UP_C4 | UP_C5 | UP_C6 | DOWN_C1 | DOWN_C2 | DOWN_C3 |
|---|---|---|---|---|---|---|---|---|---|
| ABI5 | 7.10e-04 | 0.0275 | ns | 0.034 | ns | 2.51e-05 | ns | ns | ns |
| GBF2 | 2.12e-06 | 0.00133 | 0.047 | 0.0065 | ns | 2.36e-04 | ns | ns | ns |
| GBF3 | 2.7e-08 | 0.0173 | 4.1e-05 | ns | ns | 0.00187 | ns | ns | ns |
| ABF3 | 1.54e-08 | 0.0166 | ns | 6.10e-04 | ns | 2.39e-05 | ns | ns | ns |
| ANAC32 | 1.42e-06 | 2.8e-05 | ns | 0.00247 | 0.00201 | 1.01e-07 | ns | ns | ns |
| ATAF1 | ns | 0.041 | 0.0034 | 0.049 | ns | 0.0037 | ns | ns | ns |
| HB40 | 0.0032 | 8.3e-05 | 3.8e-07 | ns | 0.009 | ns | ns | ns | 0.0123 |
| HB53 | 0.0239 | 3.50e-04 | 6.4e-05 | ns | 1.70e-04 | ns | ns | ns | ns |
| HB21 | 0.00168 | 1.59e-04 | 4,00E-04 | ns | ns | ns | ns | ns | ns |

**Supplemental Table 2. Contribution of the BTFs to the regulation of *BRC1*-dependent genes.** Significance analysis of the overlap between BTF targets and genes of each *BRC1*-dependent cluster (UP_C1 to DOWN_C3). Very significant overlaps are higlighted. ns, not significant. An hypergeometric test as implemented in the Bioconductor package SuperExactTEst (Wang et al 2015) was applied. Data obtained from BRC1NET (https://greennetwork.us.es/BRC1NET/)

| BTF/Cluster | UP_C1 | UP_C2 | UP_C3 | UP_C4 | UP_C5 | UP_C6 | DOWN_C1 | DOWN_C2 | DOWN_C3 |
|---|---|---|---|---|---|---|---|---|---|
| ABI5 | 1.51 | 1.36 | ns | 1.37 | ns | 1.63 | ns | ns | ns |
| GBF2 | 1.33 | 1.26 | 1.1 | 1.23 | ns | 1.25 | ns | ns | ns |
| GBF3 | 1.28 | 1.14 | 1.23 | ns | ns | 1.1 | ns | ns | ns |
| ABF3 | 1.37 | 1.1 | ns | 1.28 | ns | 1.27 | ns | ns | ns |
| ANAC32 | 1.43 | 1.43 | ns | 1.33 | 1.35 | 1.46 | ns | ns | ns |
| ATAF1 | ns | 1.26 | 1.37 | 1.27 | ns | 1.34 | ns | ns | ns |
| HB40 | 1.21 | 1.33 | 1.39 | ns | 1.24 | ns | ns | ns | 1.17 |
| HB53 | 1.26 | 1.51 | 1.53 | ns | 1.6 | ns | ns | ns | ns |
| HB21 | 1.46 | 1.65 | 1.56 | ns | ns | ns | ns | ns | ns |

**Supplemental Table 3. Contribution of the BTFs to the regulation of *BRC1*-dependent genes.** Enrichment analysis of BTF targets in genes of each *BRC1*-dependent cluster (UP_C1 to DOWN_C3). ns, not significant. Data obtained from BRC1NET (https://greennetwork.us.es/BRC1NET/)

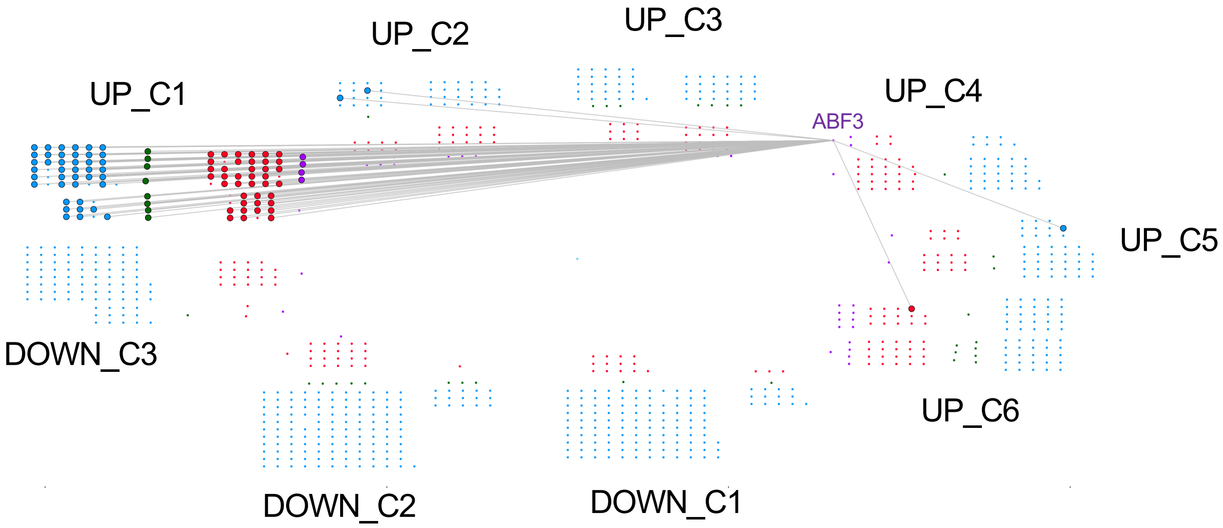


**Supplemental Fig. 7.** ABF3 targets in the BRC1 network are very significantly enriched in UP_C1 genes. Larger dots correspond to ABF3 targets in the UP_C1 cluster. Red and purple dots are BRC1 direct targets, blue and green dots are indirect BRC1 targets. Purple and green dots represent genes encoding TFs. Nodes outside UP_C1 are genes belonging to several clusters (including UP_C1). Data generated using BRC1NET (https://greennetwork.us.es/BRC1NET/)

a

| Motif Name | # occ in BRC1NET | Avg. # random occ. | SD random occ. | Estimated p-value |
|---|---|---|---|---|
| Regulated Feedback Loop | 264 | 85.89 | 16.08 | < 1e-3 |
| Feedback Loop with a single regulator* | 367 | 254.71 | 31.52 | < 1e-3 |
| Feed Forward Loop | 30185 | 25824.36 | 1420.77 | < 1e-3 |
| Feedback Loop with multiple output | 16850 | 8997.55 | 1142.18 | < 1e-3 |
| Feedback Loop with individual output** | 40752 | 26065.16 | 3104.3 | < 1e-3 |
| Double Feedback Loop*** | 188 | 86.94 | 21.71 | < 1e-3 |
| Feedback Loop with long branch*** | 151 | 65.318 | 13.82 | < 1e-3 |
| Double Feedback Loop with extra branch*** | 343 | 118.92 | 27.06 | < 1e-3 |
| Triple Feedback Loop | 70 | 13.62 | 6.35 | < 1e-3 |

* Regulated Feedback Loop variant
**Feedback Loop with multiple output variant
***Feedback Loop variant

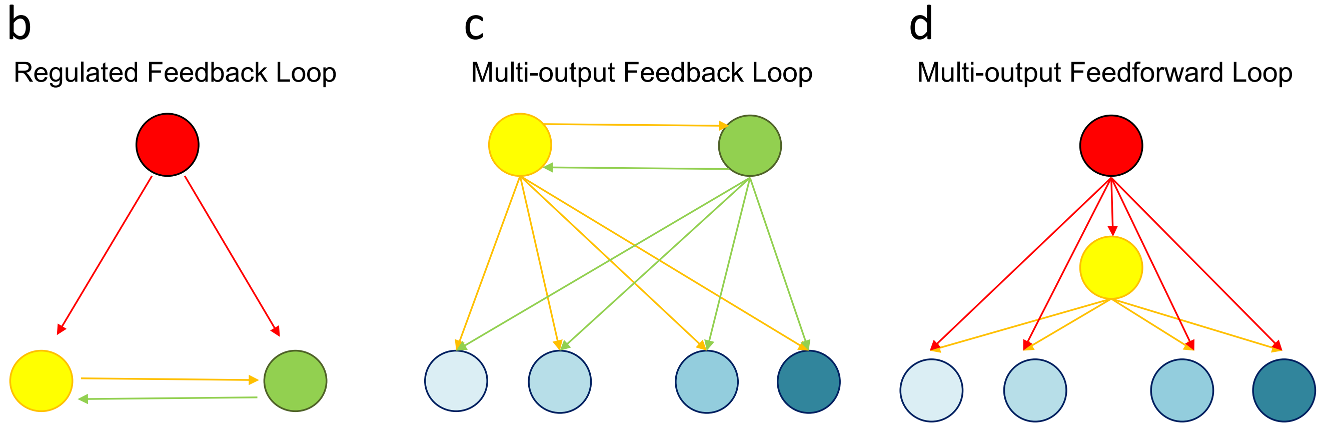


**Supplemental Fig. 8. Network motifs identified in the transcriptional network downstream of BRC1. a,** Subgraphs represent regulatory modules significantly overrepresented in the transcriptional network downstream of BRC1 when compared to random networks. These regulatory motifs are expected to play a key role in the dynamics of the network. **b,** Regulated Feedback Loop: a master regulator directly regulates two TFs that in turn regulate each other establishing a feedback loop. **c,** Multi-output Feedback Loop, two TFs regulate each other in a feedback loop to jointly regulate the same set of target genes. **d**, Multi-output Feedforward Loop, a master regulator directly regulates a set of target genes and simultaneously regulates a second TF that in turn controls the same set of target genes.

| BTFs/Cluster | UP_C1 | UP_C2 | UP_C3 | UP_C4 | UP_C5 | UP_C6 | DOWN_C1 | DOWN_C2 | DOWN_C3 |
|---|---|---|---|---|---|---|---|---|---|
| BRC1/ATAF1 | 7.01e-04 | 3.9e-04 | 0.0053 | 4.40e-04 | ns | 5.3e-09 | ns | ns | ns |
| BRC1/HB40 | 7.4e-06 | 7.3e-07 | 8.80e-04 | 0.0030 | ns | ns | ns | ns | ns |
| BRC1/WRKY18 | 0.00233 | ns | 0.0082 | 0.0113 | ns | 3.6e-05 | ns | ns | ns |
| ABI5/GBF2 | 9.3e-08 | 3.8e-04 | 0.0068 | 0.00007 | ns | 9.3e-10 | ns | ns | ns |
| ABI5/GBF3 | 1.78e-06 | 0.00305 | 0.0109 | 0.0033 | ns | 3.7e-08 | ns | ns | ns |
| ABI5/ABF3 | 1.08e-05 | 0.0099 | ns | 4.70e-04 | ns | 6.5e-08 | ns | ns | ns |
| ABI5/ANAC032 | 1.95e-04 | 3.6e-04 | ns | 8.00e-04 | ns | 5.7e-12 | ns | ns | ns |
| ABI5/ATAF1 | 6.5e-04 | 5.3e-05 | 0.0142 | 0.0061 | ns | 4.00e-08 | ns | ns | ns |
| ABI5/HB40 | 1.00e-04 | 0.0037 | 1.6e-04 | ns | ns | 0.0067 | ns | ns | ns |
| ABF3/ANAC032 | 3.7e-12 | 4.4e-10 | 0.0225 | 1.02e-09 | 3.5e-07 | 1.25e-13 | ns | ns | ns |
| ABF3/GBF2 | 1.05e-15 | 7.3e-09 | 3.6e-05 | 2.92e-10 | 0.0029 | 3.3e-14 | ns | ns | 0.0104 |
| ANAC032/GBF3 | 2.26e-11 | 5.5e-10 | 0.0106 | 1.59e-05 | 1.9e-05 | 1.92e-11 | ns | ns | ns |
| ANAC032/GBF2 | 7.7e-10 | 8.00e-10 | 0.0088 | 4.2e-08 | 0.0011 | 1.07e-10 | ns | ns | ns |

**Supplemental Table 4. Contribution of FBLs to the regulation of the *BRC1*-dependent genes.** Significance analysis of the overlap of targets of BTFs involved in FBLs with *BRC1*-dependent clusters (UP_C1 to DOWN_C3). Very significant overlaps are highlighted. ns, not significant. An hypergeometric test as implemented in the Bioconductor package SuperExactTEst (Wang et al 2015) was applied. Data generated using BRC1NET (https://greennetwork.us.es/BRC1NET/)

| BTFs/Cluster | UP_C1 | UP_C2 | UP_C3 | UP_C4 | UP_C5 | UP_C6 | DOWN_C1 | DOWN_C2 | DOWN_C3 |
|---|---|---|---|---|---|---|---|---|---|
| BRC1/ATAF1 | 1.98 | 2.2 | 1.84 | 2.3 | ns | 2.79 | ns | ns | ns |
| BRC1/HB40 | 1.95 | 2.23 | 1.73 | 1.75 | ns | ns | ns | ns | ns |
| BRC1/WRKY18 | 1.68 | ns | 1.62 | 1.69 | ns | 1.94 | ns | ns | ns |
| ABI5/ABF3 | 1.97 | 1.61 | ns | 1.94 | ns | 2.18 | ns | ns | ns |
| ABI5/GBF2 | 2.25 | 1.91 | 1.61 | 2.14 | ns | 2.41 | ns | ns | ns |
| ABI5/GBF3 | 1.96 | 1.65 | 1.5 | 1.7 | ns | 2.09 | ns | ns | ns |
| ABI5/ANAC032 | 1.95 | 2.05 | ns | 2.06 | ns | 2.84 | ns | ns | ns |
| ABI5/ATAF1 | 2.08 | 2.53 | 1.78 | 2.06 | ns | 2.83 | ns | ns | ns |
| ABI5/HB40 | 1.89 | 1.73 | 1.92 | ns | ns | 1.57 | ns | ns | ns |
| ABF3/ANAC032 | 1.99 | 2.01 | 1.31 | 2.08 | 1.92 | 2.02 | ns | ns | ns |
| ABF3/GBF2 | 1.96 | 1.79 | 1.52 | 1.93 | 1.44 | 1.88 | ns | ns | 1.28 |
| ANAC032/GBF3 | 1.84 | 1.89 | 1.32 | 1.67 | 1.68 | 1.82 | ns | ns | ns |
| ANAC032/GBF2 | 1.91 | 2.04 | 1.38 | 2.01 | 1.59 | 1.93 | ns | ns | ns |

**Supplemental Table 5. Contribution of FBLs to the regulation of the *BRC1*-dependent genes.** Enrichment of targets of BTFs involved in FBLs in genes of each cluster. Data generated using BRC1NET (https://greennetwork.us.es/BRC1NET/)

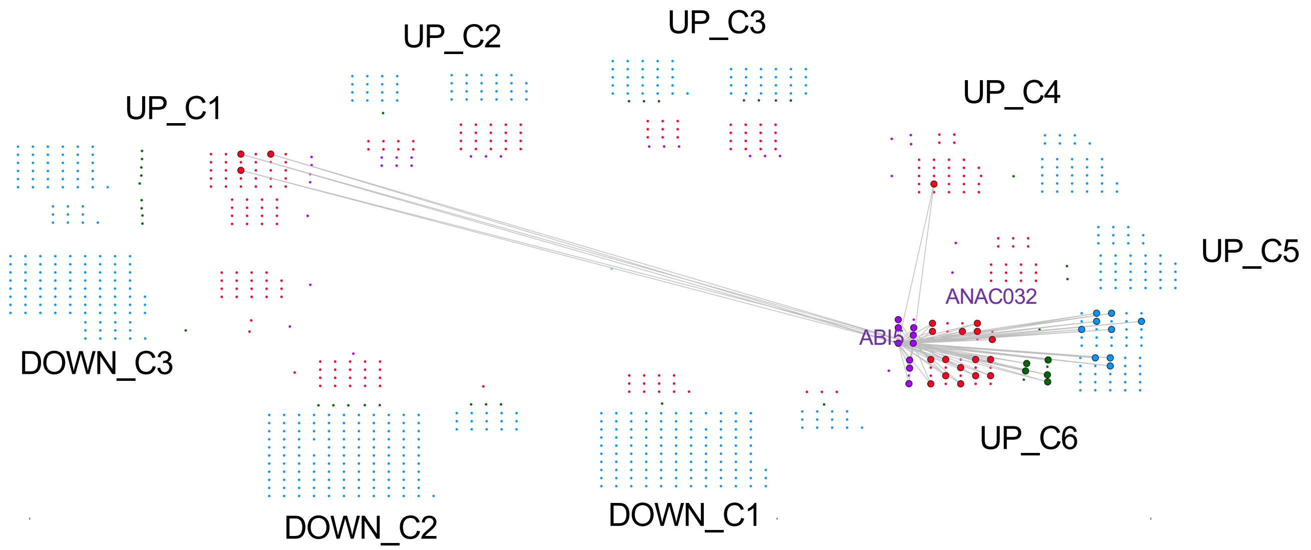


**Supplemental Fig. 9. Common targets of ABI5 and ANAC032 are significantly enriched in genes from UP_C6.** ABI5 and ANAC032 form a multi-output FBL whose common targets are indicated. Larger dots correspond to common targets of ABI5 and ANAC032. Red and purple dots are BRC1 direct targets, blue and green dots are indirect BRC1 targets. Purple and green dots represent genes encoding TFs. Those nodes outside UP_C6 are genes belonging to several clusters (including UP_C6).
Data generated using BRC1NET (https://greennetwork.us.es/BRC1NET/)

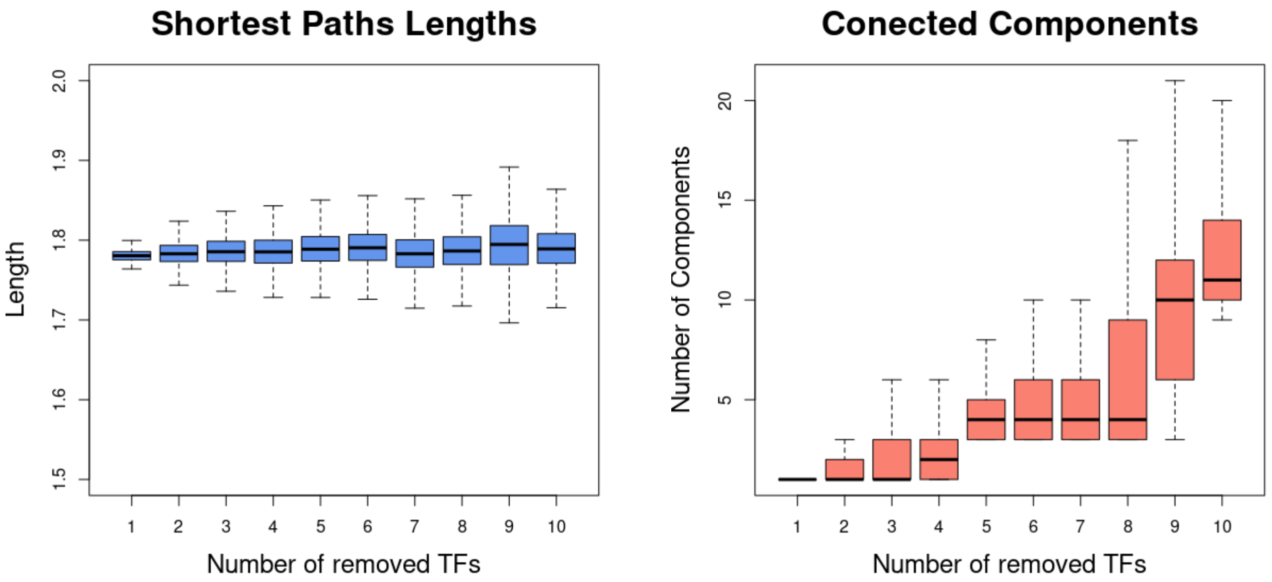


**Supplemental Fig. 10. The BRC1 network is robust against mutation**. **a,** Effect of TF removal on the shortest path lengths connecting genes in the BRC1 transcriptional network. The mean path length is very robust and exhibits similar values independently of the number of TFs removed (excepting BRC1). **b**, Effect of TF removal on the BRC1 network connectivity. Numbers of isolated connected components is represented. The network remains highly connected in spite of increasing number of removed TFs, and it shows a low number of different isolated connected components.

| NAME | SEQUENCE 5'-3' |
|---|---|
| ABI5_FW | 5'-GGGGACAAGTTTGTACAAAAAAGCAGGCTtctattcaaaatctaacaag-3' |
| ABI5_REV | 5'-GGGGACCACTTTGTACAAGAAAGCTGGGTttaacaactgcatcatatac-3' |
| ABF3_#2_F | 5'-GGGGACAAGTTTGTACAAAAAAGCAGGCTCGGGAAAGAGCACGTCTCAA-3' |
| ABF3_#2_R | 5'-GGGGACCACTTTGTACAAGAAAGCTGGGTCGTACAACTAGGAGGATCAA-3' |
| GBF2_FW2 | 5'-GGGGACAAGTTTGTACAAAAAAGCAGGCTgttatagccttaatggtttt-3' |
| GBF2_REV2 | 5'-GGGGACCACTTTGTACAAGAAAGCTGGGTtgctcaatgcagagtaatgacacag-3' |
| GBF3_FW2 | 5'-GGGGACAAGTTTGTACAAAAAAGCAGGCTtagtccaagatagaatttgggcc-3' |
| GBF3_REV | 5'-GGGGACCACTTTGTACAAGAAAGCTGGGTaggaatggtttcaaggtttccc-3' |
| MYB3_FW | 5'-GGGGACAAGTTTGTACAAAAAAGCAGGCTttgtgcatagggaaccttcg-3' |
| MYB3_REV | 5'-GGGGACCACTTTGTACAAGAAAGCTGGGTctttgttgatctctatatgt-3' |
| ATAF1_FW | 5'-GGGGACAAGTTTGTACAAAAAAGCAGGCTaatataattaacaccgtcga-3' |
| ATAF1_REV | 5'-GGGGACCACTTTGTACAAGAAAGCTGGGTagtttaccctctttcttaat-3' |
| NAP_FW | 5'-GGGGACAAGTTTGTACAAAAAAGCAGGCTttaatcctatccttttgtc-3' |
| NAP_REV | 5'-GGGGACCACTTTGTACAAGAAAGCTGGGTgattttcagacaatttagaa-3' |

**Supplemental Table 6.** Primers used to amplify the promoter regions of the genes studied.
